# Photochemical modification of two fluorene-based molecules yields structurally distinct DNA intercalators with potent anti-MRSA activity

**DOI:** 10.1101/2025.08.22.671758

**Authors:** Avery Gaudreau, Matthew D. Beckner, Vincent Du, Ronald S. Flannagan, Varsha Balaji, Evangelos Papalambropoulos, Chenfangfei Shen, Omar M. El-Halfawy, Elizabeth R. Gillies, David E. Heinrichs

## Abstract

*Staphylococcus aureus* is a leading cause of skin and soft tissue infections, endocarditis, and bloodstream infections worldwide. The emergence of methicillin-resistant *S. aureus* (MRSA) and growing resistance to last-resort antibiotics like vancomycin have created an urgent need for new antimicrobials with distinct mechanisms of action. In this study, we characterize DB10, a planar, fluorene-based compound identified in a high-throughput screen for MRSA inhibitors. Upon UVA exposure, DB10 undergoes photoconversion from a red-colored form (DB10-R) to a yellow-colored form (DB10-Y). In comparison with DB10-R, DB10-Y exhibits reduced hydrophobicity, lower cytotoxicity, and modestly improved minimum inhibitory concentrations (MICs) towards a number of Gram-positive bacteria. DB10-Y intercalates into DNA and induces double-stranded breaks, yet resistance emerged only at low levels after prolonged serial passaging. To optimize this scaffold, we screened a panel of fluorene analogs and identified the photoconverting analog DB33, which in its yellow form (DB33-Y) is non-toxic and retained DNA intercalation activity. DB33-Y was effective against intracellular *S. aureus* in macrophages and epithelial cells and significantly reduced bacterial burden and lesion size in a murine skin infection model. DB10-Y and DB33-Y both also suppressed expression of α-toxin at sub-MIC concentrations, indicating an additional anti-virulence effect. Together, these findings highlight the therapeutic potential of fluorene-based DNA intercalators as a new class of antimicrobial and anti-virulence agents against MRSA.

## INTRODUCTION

*Staphylococcus aureus* is an opportunistic pathogen of significant concern that causes a range of infections, varying from superficial skin and soft tissue infections to severe conditions such as pneumonia, osteomyelitis, and sepsis^1^. As one of the leading causes of bacteremia and endocarditis worldwide, *S. aureus* accounts for a significant proportion of nosocomial infections^1^. The substantial clinical burden of *S. aureus* is further complicated by its remarkable ability to display resistance to virtually every antibiotic introduced into clinical use^2^. In particular, methicillin-resistant *S. aureus* (MRSA) remains a leading cause of morbidity and mortality in both healthcare and community settings, and resistance has increasingly been observed against glycopeptides, lipopeptides, and linezolid, which are among the few remaining treatment options^3,4^. This persistent rise in resistance highlights an urgent need for new antimicrobial agents that target bacterial vulnerabilities beyond the scope of traditional drug targets such as cell wall synthesis, protein translation, and DNA replication^5^.

A promising class of antimicrobial agents includes compounds that target bacterial DNA through intercalation and strand damage^6–8^. DNA intercalators act by inserting planar, aromatic moieties—such as those found in naphthoquinones, anthracyclines, and fluorenes—between base pairs of the DNA helix, distorting its structure and disrupting critical processes like replication, transcription, and repair^9,10^. Several DNA intercalators, including well-studied drugs like doxorubicin, have demonstrated antimicrobial activity in addition to their anticancer properties^11,12^. These compounds often exert a multifaceted mechanism of action, not only damaging DNA directly but also generating reactive oxygen species, disrupting gyrase or topoisomerase activity, or triggering broader stress responses within bacterial cells^13,14^. Other intercalators, including those with naphthoquinone, indole, and fluorene moieties, have shown activity against MRSA, highlighting the potential of this compound class for use as anti-staphylococcals^14–18^.

Beyond their activity against planktonic cells, many DNA-targeting agents also exhibit efficacy against biofilms, likely due to their ability to bind and disrupt extracellular DNA (eDNA), a critical structural component of the biofilm matrix^10,19^. This is particularly relevant for pathogens like *S. aureus* that form resilient biofilms that contribute to chronic infections and treatment failure^20^. Furthermore, these compounds can promote the curing of resistance plasmids, either by directly damaging plasmid DNA or by interfering with replication and segregation, thereby resensitizing bacteria to antimicrobials and limiting the spread of resistance^21^. Collectively, these properties highlight DNA-intercalating compounds as promising candidates for antimicrobial development.

However, despite their potential as antimicrobials, the use of DNA-targeting agents is often limited by their toxicity to host cells. Many of these compounds lack sufficient specificity for bacterial DNA and can induce genotoxic effects in mammalian cells, limiting their clinical utility^21,22^. Moreover, DNA intercalating agents are often hydrophobic, a property that facilitates their insertion between DNA bases^23^. However, this hydrophobicity also limits clinical viability due to poor solubility and a tendency to precipitate out of aqueous solutions^24^. Additionally, DNA intercalation may promote resistance in the long term. Specifically, DNA damage can activate the bacterial SOS response, a global stress pathway that enhances mutagenesis through error-prone DNA polymerases, potentially accelerating the emergence of resistant phenotypes, not only to the DNA-targeting agent itself but also to unrelated antibiotics^25,26^. Nevertheless, the benefits of DNA intercalators can outweigh these limitations, many of which can be addressed through rational structural modifications following the identification of a suitable lead compound.

Here, we define the mechanism of action and spectrum of activity of a planar, fluorene-containing compound named DB10, identified serendipitously from a high-throughput screen for inhibitors active against MRSA. Interestingly, upon UVA exposure, DB10 photoconverted from a red-colored compound (DB10-R) to a yellow-colored compound (DB10-Y). The corresponding structural change modestly lowered the MIC across most bacterial isolates tested but significantly reduced the hydrophobicity and toxicity of the compound. Given these improved properties, DB10-Y was pursued as the lead compound. As predicted based on its fluorene core, DB10-Y intercalated into DNA and induced structural damage, resulting in double-stranded DNA breaks. Attempts to generate resistance to B10-Y resulted in only low-level resistance after 46 days of serial passaging in the presence of sub-MIC compound. To examine key structural features of this molecule class, a series of fluorene analogs were obtained and, of these, only one, DB33, retained strong antimicrobial activity. Like DB10, DB33 intercalated into DNA, but notably, no toxicity was observed. In addition, DB33 was effective against intracellular *S. aureus* reservoirs and reduced bacterial burden and lesion size in a murine skin infection model. Furthermore, at sub-MIC concentrations we observed a decrease in production of *S. aureus* alpha toxin (encoded by the *hla* gene), indicating an anti-virulence effect. Together, these findings highlight the therapeutic potential of the DB10 scaffold and underscore the broader utility of planar aromatic intercalators as a class of antimicrobials.

## RESULTS

### High-throughput screen identifies a photoconvertible inhibitor of MRSA

A high-throughput screen of the Maybridge library of bioactive compounds was previously conducted^27,28^ to identify potent inhibitors of *Staphylococcus aureus* USA300 LAC, the predominant community-acquired MRSA clone in North America. This initial screen, performed in tryptic soy broth (TSB), yielded 936 potential inhibitors, though early efforts focused exclusively on compounds suspected to disrupt metal ion homeostasis. More recently, we revisited this set of bioactives to identify additional promising scaffolds that showed anti-staphylococcal activity.

During this analysis, we observed that roughly 11% of the 936 hits shared a common feature in that they possessed planar, aromatic structures, chemical motifs either known to, or with the possibility to, intercalate into DNA. DNA intercalating agents may disrupt essential bacterial processes such as replication and transcription, making them attractive candidates for antimicrobial development despite potential concerns about genotoxicity^21,22^. However, when this set of 936 initial hits were tested in a Chelex-100–treated, chemically defined medium containing glucose (CDM-G), only three of these planar compounds retained their activity against USA300 LAC. The Chelex treatment removes divalent metal ions, creating a metal-depleted, host-relevant environment where some metal-dependent compounds may lose efficacy, helping to identify inhibitors with robust, metal-independent activity^28,29^. The three active compounds included one with an indole moiety, one with a phenanthroline, and one with a fluorene. Given the relative prevalence of fluorene-containing compounds (∼10%) among the sub-group of hits containing a planar aromatic moiety, and the robust activity of this compound under metal-depleted conditions, we selected it for further study and designated it DB10 (see **Figure 1A** for structure).

**Figure 1:**
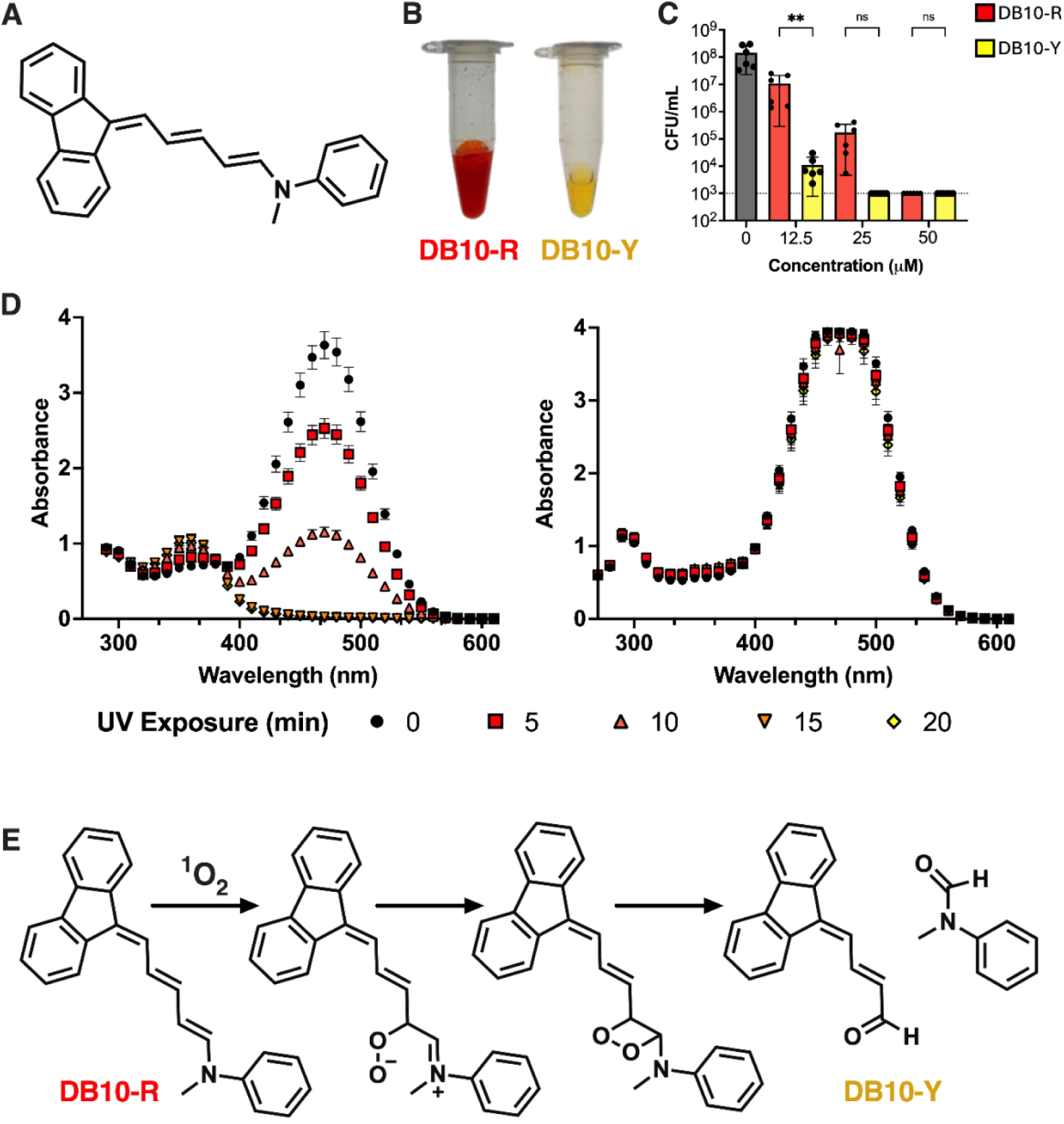
DB10 is a photoconvertible inhibitor of USA300 LAC. **(A)** Chemical structure of DB10. In **(B)** the image depicts aliquots of DB10 before (DB10-R) and after (DB10-Y) photoconversion. (**C**) Killing of *S. aureus* following 24 h exposure to DB10-R or DB10-Y at the indicated concentrations. Bacteria were normalized to OD₆₀₀ = 1 in PBS, treated with compound or vehicle, and incubated at 37°C for 24 h before serial dilution and plating on TSA to enumerate CFU. Data are shown as the mean ± SD from at least three biological replicates. ∗∗p ≤ 0.01 using a one-way ANOVA with Dunnet’s multiple comparison. **(D)** Change in absorbance spectra of DB10-R as it is converted to DB10-Y during long wave UV (UVA, 365nm) or (**E**) short wave UV (UVB, 255nm) exposure. Absorbance was read every 5 mins and data are shown as the mean ± SD from at least three independent experiments. In (**F)** ^1^H NMR-spectroscopy was used to predict the depicted conversion pathway of DB10-R to DB10-Y upon UV-A exposure.

Interestingly, during our initial experiments using DB10, we observed a gradual color change from red (DB10-R) to yellow (DB10-Y) after the compound was reconstituted and exposed to glass-filtered sunlight over a period of 3–5 days (**Figure 1B**). This observation suggested that DB10 may undergo photoconversion, a phenomenon in which light exposure induces a chemical change in a compound and also potentially alters its biological activity or stability. Indeed, photoconversion of DB10-R to DB10-Y altered the bactericidal activity of these compounds against *S. aureus*. While both forms exhibit similar efficacy at 50 μM, DB10-Y consistently results in lower CFU/mL counts at 12.5 and 25 μM, suggesting that the yellow form is more anti-bacterial (**Figure 1C**). Photoconversion also improved the solubility of the compound and while DB10-R precipitated from aqueous solution at concentrations above 300 μM, DB10-Y remained soluble up to 1 mM.

Next, we sought to investigate the mechanism behind this suspected photoconversion. To this end DB10 was subjected to either longwave (UVA) or shortwave (UVB) ultraviolet light, and the resulting color change was monitored over time by absorbance (**Figure 1D** and **1E**). Upon UVA exposure, a time-dependent decrease in absorbance at ∼470 nm was observed, corresponding to the disappearance of the red-colored species (DB10-R). Concurrently, the absorbance around the ∼370 nm peak increased, consistent with the formation of the yellow photoproduct (DB10-Y). Complete photoconversion of DB10-R occurred within 15 minutes of UVA exposure. In contrast, when the same concentration of DB10-R was exposed to UVB, no appreciable color change was observed, and the absorbance peak at ∼470 nm remained stable and did not decrease. This strongly suggested that UVB exposure was insufficient to induce any chemical transformation of DB10, at least under the tested conditions, and that the photoconversion of DB10-R to DB10-Y was driven by UVA.

To assess the structural basis of this conversion, ^1^H NMR analyses were performed. The appearance of an additional peak at 9.8 ppm confirmed a structural change (**Supplementary Figures S1-S3**), indicating the presence of a new aldehyde. The predicted conversion pathway and resulting structure of DB10-Y are shown in **Figure 1F**.

### Photoconversion enhances activity of DB10 against Gram-positive bacteria

To further assess the spectrum of activity and bacterial response to both DB10-R and DB10-Y, we determined the minimum inhibitory concentrations (MICs) of each compound against various *S. aureus* strains, other *Staphylococcus* species, and non-staphylococcal bacteria. In Mueller–Hinton Broth (MHB), both DB10-R and DB10-Y inhibited the majority of *Staphylococcus* isolates tested (**Table 1**), as well as other Gram-positive species at concentrations ≤ 50 µM. However, at the highest concentration tested (100 µM), growth persisted for *S. cohnii*, *S. warneri*, *B. subtilis*, and *S. pyogenes* in the presence of DB10-R. Neither form of the compound at the concentrations tested here showed activity against the Gram-negative species evaluated (**Table 1**). Although not statistically significant for most isolates, and consistent with CFU counts, DB10-Y consistently exhibited slightly lower MIC values than DB10-R, suggesting a modest increase in potency following photoconversion.

**Table 1:** DB10-R and DB10-Y are potent inhibitors of Gram-positive bacteria. MICs of select (**A**) *S. aureus* strains, (**B**) Staphylococcal spp. and (**C**) non-staphylococcal spp. The concentration of DB10-R or -Y that completely inhibited bacterial growth was determined to be the MIC. At least three biological replicates for each isolate and form of DB10 were conducted to determine the MICs.

| Bacterial isolate | DB10-R (μM) | DB10-Y (μM) |
| --- | --- | --- |
| *S. aureus* USA100 | 50 | 25 |
| *S. aureus* USA200 | 50 | 25 |
| *S. aureus* USA300 | 50 | 25 |
| *S. aureus* USA400 | 50 | 50 |
| *S. aureus* USA600 | 100 | 100 |
| *S. capitis* | 50 | 25 |
| *S. epidermidis* | 50 | 25 |
| *S. chromogenes* | 50 | 25 |
| *S. lugdunensis* | 50 | 50 |
| *S. cohnii* | >100 | >100 |
| *S. warneri* | >100 | 50 |
| *B. subtilis* | >100 | 100 |
| *S. pyogenes* | 100 | 50 |
| *S. agalactiae* | 50 | 50 |
| *E. faecalis* | 25 | 25 |
| *P. aeruginosa* | >100 | >100 |
| *E. coli* | >100 | >100 |

### *S. aureus* alters its transcriptome upon exposure to DB10-R and DB10-Y

To understand how *S. aureus* responds to each compound and whether DB10-R and DB10-Y evoked conserved transcriptional responses, we performed RNA-seq on *S. aureus* USA300 LAC cultures exposed to sub-MIC concentrations of each compound for one hour during early exponential phase. Although MICs in **Table 1** were determined in Mueller-Hinton broth (MHB), the standard medium for antimicrobial testing, *S. aureus* is more typically grown in tryptic soy broth (TSB), which is also the conventional medium for transcriptomic analyses. Notably, the MICs for both DB10-R and DB10-Y in TSB are approximately 100 µM. Untreated cultures were included as controls to establish baseline gene expression. A total of 59 and 71 differentially expressed genes (DEGs) were identified following exposure to DB10-R (**Figure 2A**) and DB10-Y (**Figure 2B**), respectively.

**Figure 2:**
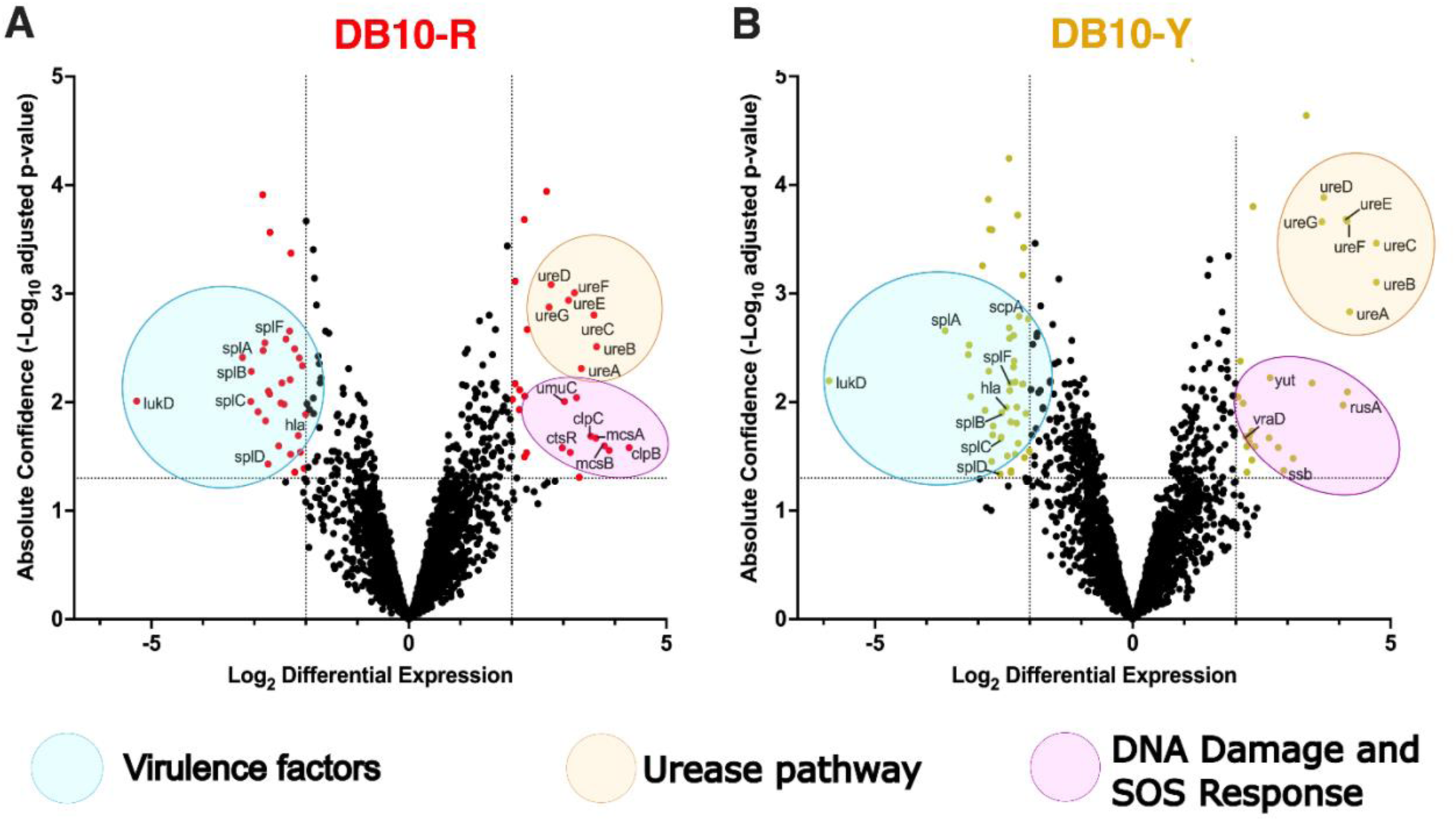
DB10-R and DB10-Y elicit similar transcriptional responses. Volcano plots of RNA-seq analysis of *S. aureus* USA300 LAC after exposure to 50 μM (i.e. sub-MIC in TSB) of either (**A**) DB10-R or (**B**) DB10-Y for 1 h during early exponential phase. Results are from three independent experiments. Differentially expressed genes (absolute confidence >2, |log2 differential expression| >1.3) are colored in red for DB10-R and yellow for DB10-Y. Differentially expressed pathway groups are encircled in blue (virulence factors), orange (urease pathway) or purple (DNA damage and SOS response). Detailed RNAseq data can be found in Supplementary **Table S4** (RNAseq for *S. aureus* exposure to DB10-R) and **Table S5** (RNAseq for *S. aureus* exposure to DB10-Y).

DB10-R and DB10-Y both induced expression of genes associated with stress and DNA damage responses. Additionally, DB10-R treatment led to significant upregulation of *clpB* and *clpC*, which encode Clp family proteases involved in refolding or degrading misfolded proteins under stress conditions^30^. DB10-R also increased expression of *umuC*, *ctsR*, *mcsA*, and *mcsB*, while DB10-Y upregulated *rusA* and *ssb*. In *S. aureus*, *umuC* is part of the SOS response and encodes a DNA polymerase that bypasses damaged DNA^31^. *ctsR* regulates heat shock genes, while *mcsA* and *mcsB* are involved in controlling Clp-dependent proteostasis under stress^32^. *rusA*, although better characterized in *E. coli,* likely functions similarly in resolving Holliday junctions^33^, and *ssb* stabilizes single-stranded DNA during replication and repair^34^. Collectively, this suggests that both forms activate cellular pathways involved in genome maintenance and repair. Interestingly, both treatments also resulted in upregulation of the *ureABCDEF* operon, which encodes urease and associated accessory proteins^35,36^. Urease catalyzes the hydrolysis of urea into ammonia and carbon dioxide and plays a known role in neutralizing intracellular pH during acid stress^36^. Upregulation of this operon suggests a broader cellular response to intracellular acidification, metal dysregulation, or other stress-related cues triggered by DB10 exposure. Notably, both forms of DB10 significantly downregulated numerous virulence factors, including the *spl* serine protease-like operon, *hla* (encoding α-hemolysin), and *lukD* (a component of the leukocidin cytotoxin) (**Figure 2**).

Collectively, these data demonstrate that DB10-Y elicits similar transcriptional changes on *S. aureus* as DB10-R. Based on that, and our findings that DB10-Y also has improved solubility and modestly enhanced potency, see preceding section, we therefore selected DB10-Y for further studies.

### DB10-Y intercalates into DNA and causes dsDNA breaks

As mentioned previously, DB10-Y is a planar, aromatic compound with a core fluorene moiety. Considering other compounds with this moiety are known to either intercalate or interact with DNA in some way^37,38^, along with the induction of the DNA damage response as observed in our RNA-seq after DB10-Y exposure, we speculated that the main mechanism of action of this inhibitor would involve DNA binding in some manner.

To evaluate the DNA intercalative properties of DB10-Y, we performed a competitive intercalation assay using ethidium bromide (EtBr), a well-established DNA intercalator that fluoresces upon binding to DNA^39^. In this assay, EtBr is allowed to intercalate into DNA, and the subsequent addition of compounds that compete for intercalation sites displace EtBr, resulting in a measurable reduction in fluorescence^15^. As expected, increasing concentrations of DB10-Y resulted in a dose-dependent decrease in EtBr fluorescence, indicating that DB10-Y competitively displaces EtBr (**Figure 3A**). Notably, DB10-Y alone—whether in the presence or absence of DNA—did not emit detectable fluorescence under the assay conditions, confirming that the observed signal changes were attributable solely to EtBr displacement. To confirm the specificity of this effect, we included erythromycin, a non-intercalative control compound. Erythromycin did not alter EtBr-DNA fluorescence, reinforcing our conclusion that the fluorescence decrease observed with DB10-Y was due to its specific displacement of EtBr from DNA.

**Figure 3:**
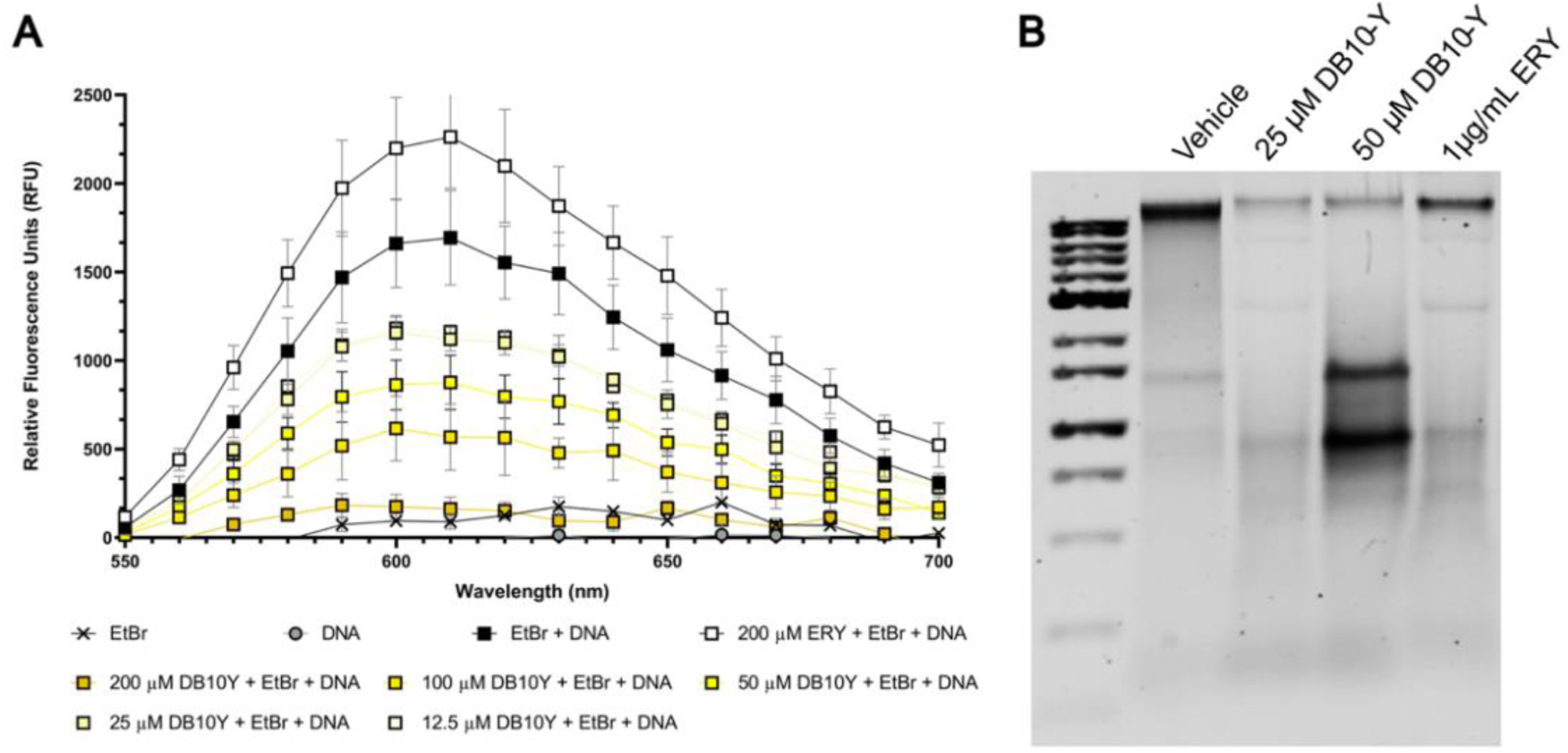
DB10-Y competitively intercalates into and causes damage to DNA. (**A**) DB10-Y displaces EtBr from DNA. EtBr, DNA and various concentrations of DB10-Y or the control erythromycin were incubated together for 30 minutes in the dark before the fluorescence spectra of EtBr was read (excitation 525 nm). DB10-Y, EtBr and DNA alone showed no intrinsic fluorescence, and ERY, a non-intercalating control, had no effect on EtBr fluorescence. Data are shown as the mean ± SD from at least three biological replicates. (**B**) Representative agarose gel of genomic DNA extracted from DB10-Y–treated cultures under neutral conditions. USA300 was subcultured overnight with varying concentrations of DB10-Y, and the genomic DNA of each sample was isolated via phenol-chloroform extraction. Samples were run on a 0.8% agarose TBE gel at 100V. Accumulation of shorter DNA fragments is indicative of double-stranded breaks (DSBs). Figure is representative from three independent replicates.

Given that intercalating agents can induce DNA damage, we next assessed whether DB10-Y caused DNA strand breaks. Genomic DNA was isolated from cultures exposed to DB10-Y and subjected to electrophoresis under either neutral or alkaline conditions to differentiate between double-stranded breaks (DSBs) and single-stranded breaks (SSBs), respectively, as described previously^40^. Under neutral conditions (pH 7.5), DB10-Y treatment resulted in the accumulation of shorter DNA fragments, consistent with the presence of DSBs (**Figure 3B**). In contrast, the DNA banding pattern under alkaline conditions was similar to that observed under neutral conditions, indicating no detectable increase in SSBs (**Figure S4A**). These findings suggest that DB10-Y induces DNA damage specifically in the form of DSBs.

Reactive oxygen species (ROS), including superoxide, have been implicated in mediating DNA damage in response to bactericidal antibiotics^41^. To evaluate whether oxidative stress contributes to the DNA damage observed, we assessed the production of superoxide using the nitroblue tetrazolium (NBT) assay, which detects superoxide through the reduction of NBT to a blue formazan precipitate^42^. Treatment of *S. aureus* cells with DB10-Y did not result in detectable ROS production, as no significant superoxide levels were observed (**Figure S4B**), suggesting that the observed DNA damage is a direct effect of DB10-Y rather than a consequence of oxidative stress.

Despite evidence of DNA intercalation and DSBs, transcriptomic analysis revealed no upregulation of key DNA damage or stress response genes, including *recA* and *rexAB* (**Figure 2B**). Furthermore, transposon mutants of these genes did not exhibit altered sensitivity to DB10-Y (**Figure S4C**). Together, these results suggest that while the primary MOA of DB10-Y involves DNA intercalation and DSB-induced damage, additional mechanisms likely contribute to its antibacterial effect.

### Mutations in *clpX* confer low-level resistance to DB10-Y

A well-established strategy for investigating antimicrobial mechanisms of action is the selection of resistant mutants through prolonged exposure. To this end, we sought to evolve resistance to DB10-Y by serially passaging cultures of USA300 LAC in the presence of sub-MIC (50 μM in TSB). After approximately 46 days, we obtained low-level but stable resistance to DB10-Y (**Figure 4A**). The extended time required to develop resistance may further suggest a pleiotropic mechanism of action that is not solely dependent on DNA intercalation and damage.

**Figure 4:**
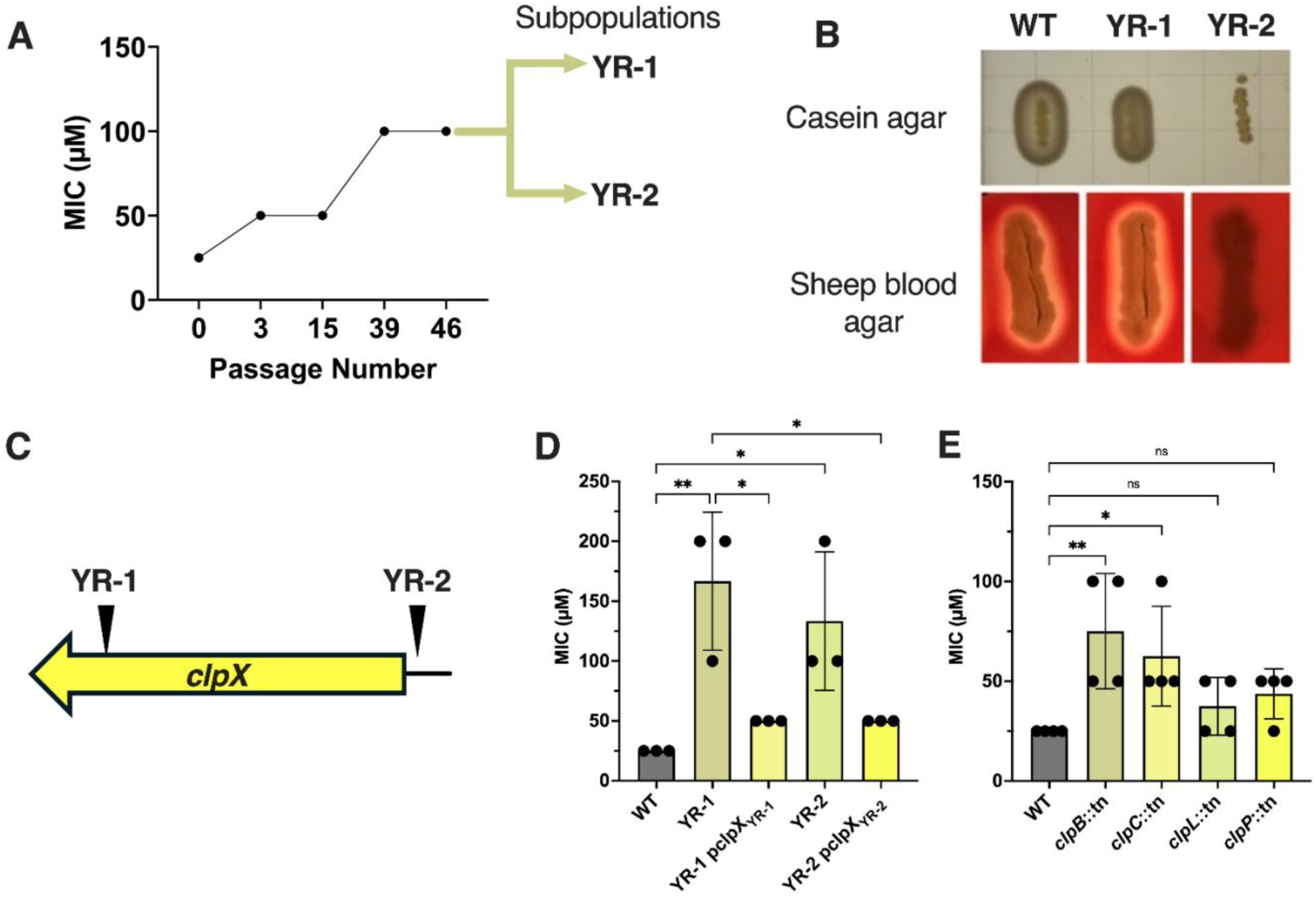
SNPs in *clpX* confer low-level resistance to DB10-Y. **(A)** Resistance development timeline of USA300 LAC exposed to sub-MIC (50 µM in TSB). Resistance was checked periodically until an increase in resistance was observed (day 39) at which time additional passaging for 7 days was done to ensure mutation stability. **(B)** Resistant sub-populations can be distinguished by a SNP in *agrC* in YR-2 resulting in a loss of casein hydrolysis and hemolysis on sheep blood agar. (**C)** Location of SNPs found in and upstream of *clpX* within the two resistant sub-populations. MICs of **(D)** resistant sub-populations overexpressing mutant *clpX* and **(E)** *clp* system transposon mutants. Data are shown as the mean ± SD of at least three independent experiments. ∗p ≤ 0.05, ∗∗p ≤ 0.01, using a one-way ANOVA with Dunnett’s multiple comparison.

Interestingly, the resistant culture consisted of two distinct subpopulations, from which we isolated representative strains YR-1 and YR-2. Subpopulation 2 (YR-2) harbored a single nucleotide polymorphism (SNP) in *agrC*, which resulted in a loss of casein hydrolysis and hemolytic activity on blood agar (**Figure 4B**), allowing it to be readily distinguished from subpopulation 1 (YR-1). Aside from the SNP in *agrC*, both subpopulations were found to contain unique SNPs within the *clpX* locus (**Figure 4C**), which encodes an ATP-dependent chaperone that interacts with ClpP to facilitate protein degradation and regulate stress responses^43^. YR-1 harboured a deletion immediately upstream of the +1 transcriptional start site of *clpX*, potentially affecting promoter function and gene expression. YR-2 contained a frameshift-inducing deletion within codon 400, near the C-terminus of *clpX*, leading to a premature stop codon. These findings suggest that alterations in *clpX* function could contribute to low-level resistance to DB10-Y.

To investigate this further, the mutant *clpX* alleles were cloned and overexpressed in either YR-1 or YR-2 backgrounds. Rather than further increasing resistance, overexpression of these alleles restored susceptibility to near wild-type levels (**Figure 4D**). To determine whether the increased resistance observed in YR-1 and YR-2, and the subsequent loss of resistance after *clpX* overexpression, was specific to DB10-Y, we compared the MICs of these mutants to wild-type USA300 LAC across other antibiotics. YR-1 displayed increased susceptibility to ofloxacin and vancomycin, while YR-2 showed increased resistance to ampicillin. In both cases, complementation with the corresponding *clpX* allele reversed the phenotype, returning susceptibility to levels comparable to USA300 LAC (**Figure S5**). These data suggest that altered *clpX* expression or activity modulates susceptibility not only to DB10-Y but also to other antibiotics, likely through effects on proteostasis or stress response pathways.

To assess whether disruption of other Clp-associated genes similarly affects susceptibility, we screened transposon insertion mutants for changes in DB10-Y resistance (**Figure 4E**). *clpB* and *clpC* transposon mutants exhibited increased resistance, while insertions in *clpP* and *clpL* did not alter susceptibility. A *clpX* transposon mutant was not available in the NTML, likely because *clpX* is essential under these conditions, preventing direct comparison. These results further suggest that perturbation of Clp-mediated proteostasis can influence susceptibility to DB10-Y. However, the low level of resistance observed, and the extended time required for emergence of this resistance indicate that these mutations likely represent supplementary, off-target adaptations rather than primary resistance mechanisms.

### Testing DB10-Y analogs identifies the less cytotoxic DB33-Y

While we initially hypothesized that the central fluorene moiety in DB10-Y was responsible for its ability to intercalate into DNA, previous studies have shown that fluorene alone is not capable of this function^44^. To identify structural features critical for antibacterial activity, we tested seven analogs of DB10, each retaining the fluorene core but with modified side chains (**Figure S6**). Analogs were selected from the Mcule database using the structure similarity search feature. Of these, six analogs exhibited no inhibitory activity against USA300 LAC at concentrations up to 400 μM (**Figure 5A**). However, one analog, designated DB33 (**Figure 5B**), displayed potent activity with an MIC of 25 μM, comparable to that of DB10.

**Figure 5:**
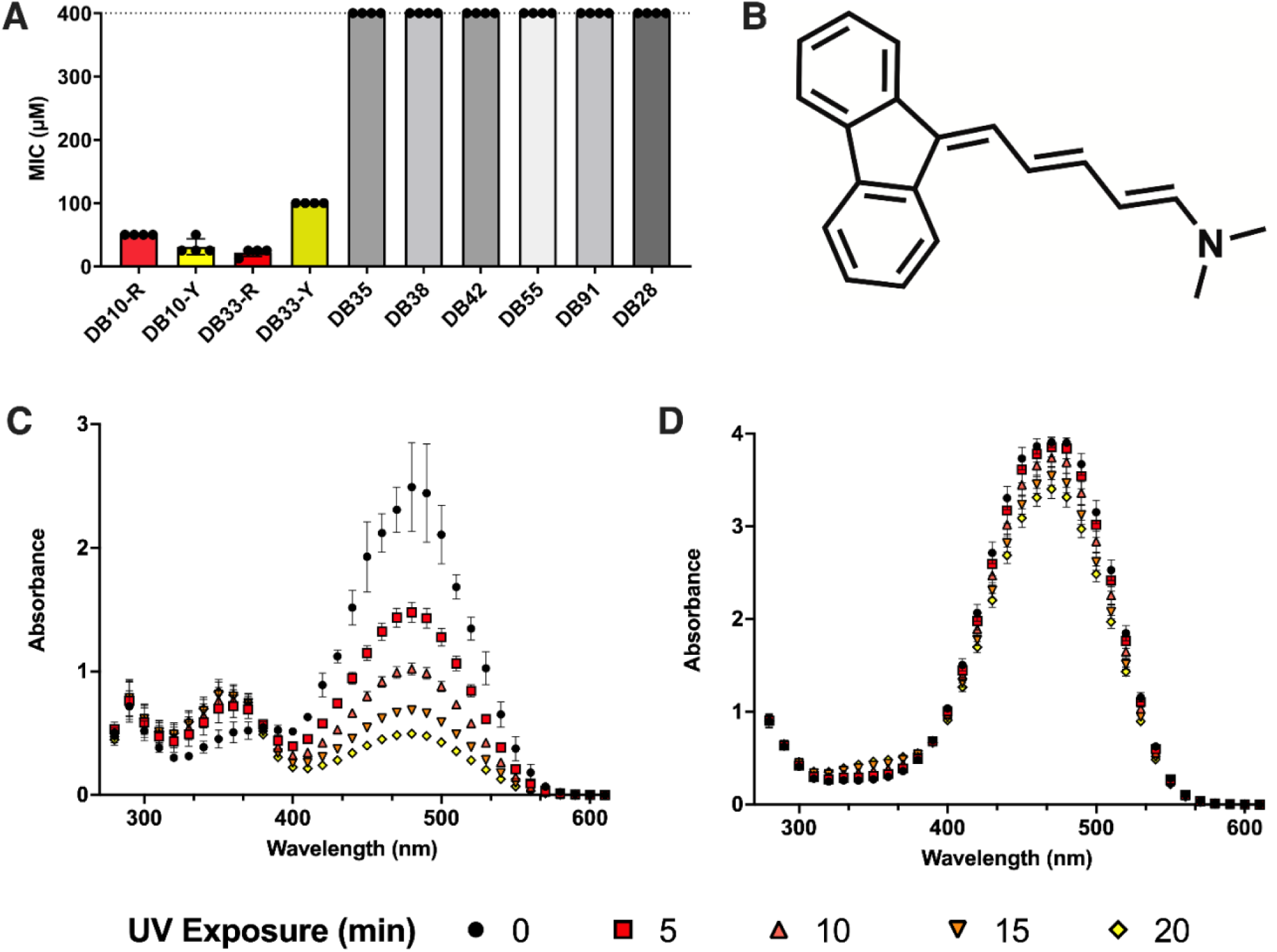
DB33 is a promising structural analog of DB10. (**A**) MICs of DB10-R and -Y with seven selected structural analogs. Analogs with no biological activity are coloured in shades of grey. (**B**) Structure of the only effective analog, DB33. (**C**) UVA-induced photoconversion of DB33-R to DB33-Y. Absorbance was read every 5 mins and data are shown as the mean ± SD from at least three replicates.

DB33, like DB10, undergoes UVA-induced photoconversion, resulting in a visible color change from red to yellow and a corresponding shift in absorbance peak from 480 nm to 350 nm (**Figure 5C**). The structure of DB10-R was confirmed by ¹H NMR (**Figure S7**). To investigate structural changes associated with the photoconversion of DB33-R, we performed ¹H NMR on DB33 at 9 g/L in DMSO-d₆. Photoconversion at this higher concentration was slower, likely due to poor light penetration. After 15 hours of irradiation at 365 nm, large amounts of starting material remained, as evidenced by a persistent alkene proton peak at 5.45 ppm (**Figure S8**). Surprisingly, switching to 450 nm light—intended to better match the absorption spectrum of DB33-R—did not improve conversion. Given the known reactivity of polyene systems with singlet oxygen^45–47^, we reasoned that oxygen depletion might be limiting conversion. Bubbling air into the solution during continued 365 nm irradiation led to full conversion after 31 hours (**Figure S9**), consistent with an oxygen-dependent mechanism.

Interestingly the ¹H NMR analysis of DB33 in **Figure S8** revealed multiple degradation products, including aldehyde peaks near 10 ppm and methyl peaks between 2.7–3.2 ppm, consistent with chain shortening and oxidation. Additionally, the disappearance of the 5.45 ppm alkene proton and downfield shifts indicate cleavage and oxidation of the polyene system. HPLC analysis of the final converted mixture after 31 hours of irradiation showed the presence of multiple peaks (**Figure S10**), further demonstrating that multiple degradation products are present.

Based on prior studies of polyene photosensitizers, two singlet oxygen–mediated degradation mechanisms are proposed: (i) stepwise reduction of the polyene system causing progressive chain shortening^45–47^ (**Figure 6A**), and (ii) oxidative cleavage producing aldehydes and other oxidized fragments^47–49^ (**Figure 6B**). A combination of these mechanisms (i.e., chain shortening followed by oxidative cleavage) can lead to a number of potential degradation products (**Figure 6C, compounds 2-7**). Other oxidative degradation products such as alcohols, epoxides, and endoperoxides may also be possible^49^, arising directly from DB33 or from the above-mentioned species (**Figure 6C, compounds 8-11**). The numerous possible cleavage and oxidation sites can account for the large number of observed products.

**Figure 6:**
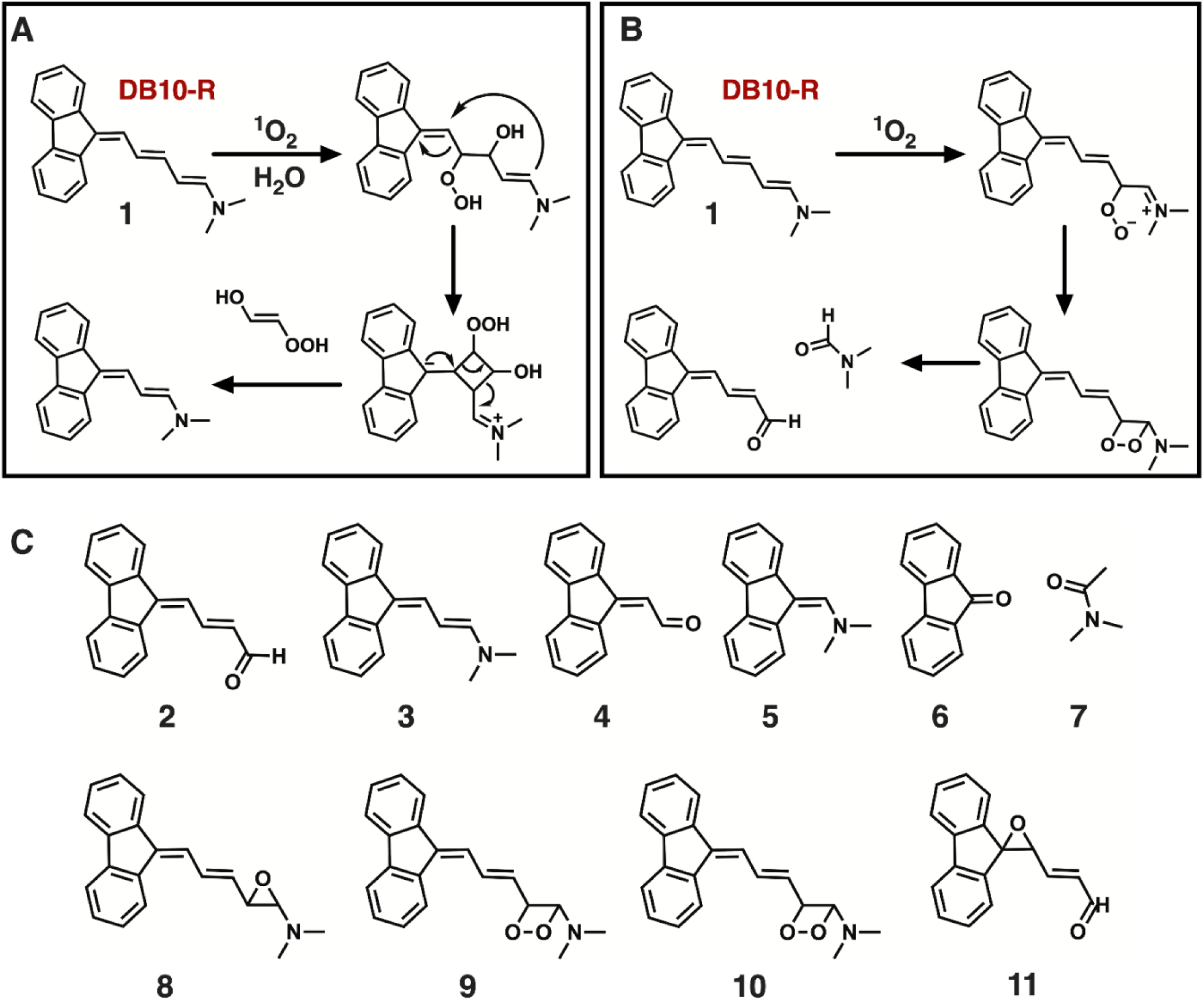
Proposed photoconversion mechanism and photo-products of DB33. Potential mechanisms of photodegradation of compound 1 (DB33), leading to (**A**) reduction in the polyene chain length and (**B**) oxidative cleavage to generate the aldehyde. (**C**) Potential photodegradation products for compound 1, arising from oxidative cleavage and polyene chain length reduction induced by singlet oxygen (compounds 2-7) and the oxidative degradation products derived from compounds 2-7 (compounds 8-11).

Mass spectrometry of the three largest HPLC peaks (3, 5, and 8) again confirmed the presence of multiple species. Peak 3 (**Figure S11**) showed m/z values at 107 and 191, the latter likely representing a sodium adduct of a dimeric fragment. Peaks 5 (**Figure S12**) and 8 (**Figure S13**) contained masses consistent with compound 2 and other oxidized derivatives of DB33. This diversity of products supports the proposed oxidative cleavage and chain shortening mechanisms. For our purposes, we will refer to this collection of degradation products simply as DB33-Y.

Notably, both the red (DB33-R) and yellow (DB33-Y) forms showed similar antibacterial activity to DB10 against a panel of *S. aureus* strains and other species (**Table S1**). Interestingly, photoconversion of DB33 resulted in a modest increase in MIC (100 μM) against USA300 LAC, opposite to the trend observed for DB10.

To confirm whether DB33-Y shares the same DNA-targeting mechanism as DB10-Y, we once again performed a competitive EtBr displacement assay. As DB33-Y concentration increased, EtBr fluorescence decreased in a dose-dependent manner, indicating that DB33-Y competes with EtBr for DNA binding and intercalates into DNA (**Figure S14A**). Consistent with this concept of a shared mechanism, *clpX* mutant strains YR-1 and YR-2, originally selected for resistance to DB10-Y, also displayed resistance to DB33-Y, further supporting that the analog DB33-Y shares a common mode of action to DB10-Y (**Figure S14B**).

We next assessed whether DB10 analogs that did not have appreciable anti-staphylococcal activity (colored in grey in Figure 5E) had the ability to intercalate into DNA by performing the EtBr displacement assay. All exhibited some degree of DNA binding ability (**Figure S15**), EtBr displacement was generally lower than that observed with DB10-Y. Interestingly, DB35 showed slightly greater displacement than DB10-Y, despite lacking antibacterial activity, indicating that strong intercalation alone may not be sufficient for antimicrobial efficacy.

### DB33-Y effectively clears intracellular bacterial reservoirs

*S. aureus* is a pathogen capable of persisting inside host cells (e.g. macrophages, dendritic cells, and endothelial cells) which can protect the bacteria from immune responses and many antibiotics^50–52^. Therefore, we sought to determine whether DB10 or DB33 compounds could be used to antagonize *S. aureus* in cellular models of infection^28,53–58^. As a first step towards this, we evaluated the cytotoxicity of these compounds using the lactate dehydrogenase (LDH) release assay, which measures LDH released from damaged mammalian cells as an indicator of membrane integrity and cellular toxicity^59^. Released LDH converts lactate to pyruvate, generating NADH, which reduces a tetrazolium salt to produce the colored formazan product^59^. Formazan production after DB10 or DB33 exposure was quantified spectrophotometrically at 500 nm. Notably, despite similar antibacterial activity and a shared intercalative mechanism, DB33-Y demonstrated significantly reduced cytotoxicity compared to both forms of DB10 (**Figure 7A**). DB33-Y caused less LDH release than DB33-R at the same concentration, and substantially less than either DB10 form, highlighting its improved safety profile.

**Figure 7:**
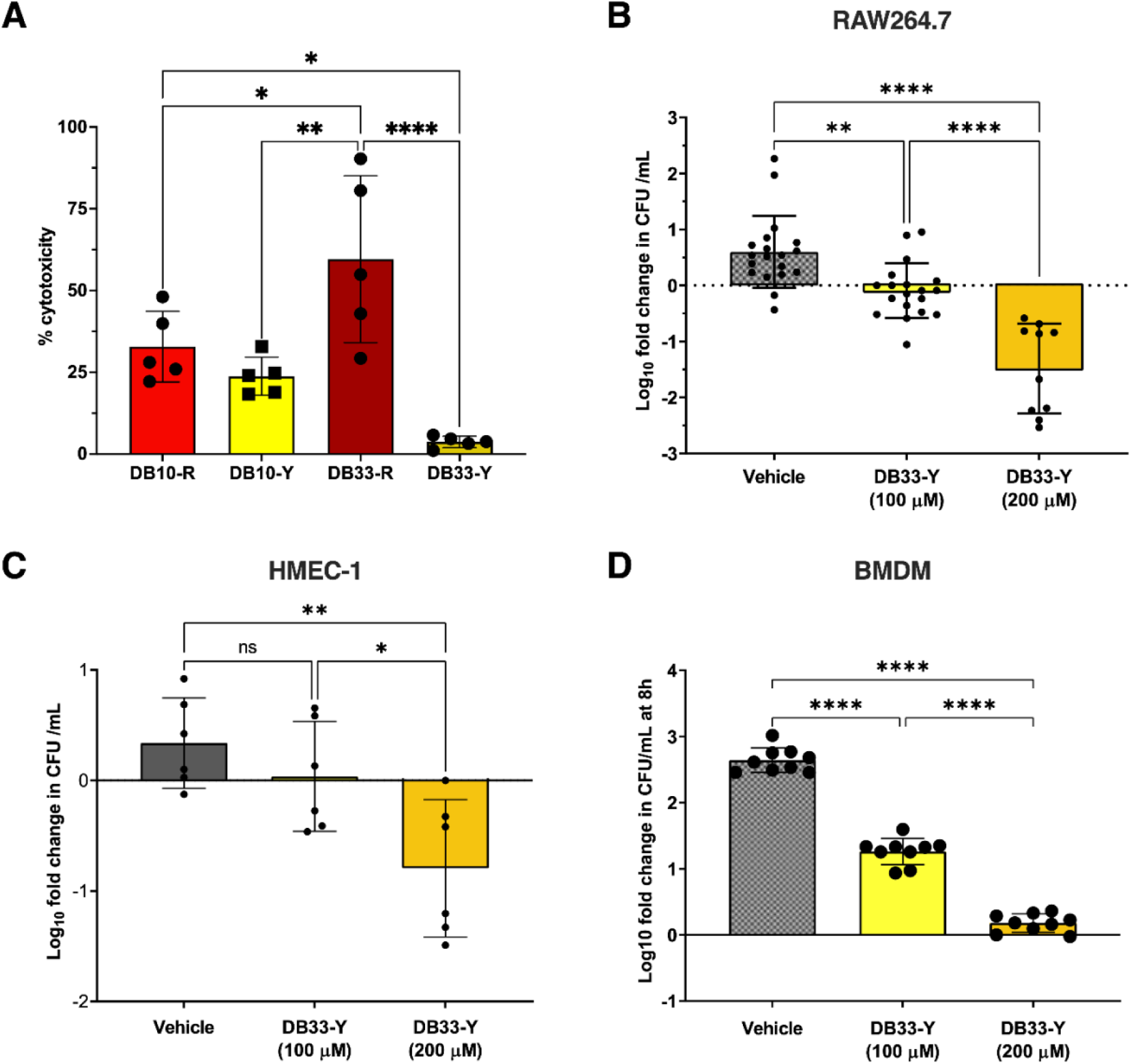
DB33-Y effectively clears intracellular bacterial reservoirs and reduces lesion sizes in a skin infection model. **(A)** Cytotoxicity of DB10 and DB33 at 100 μM in RAW264.7 cells, measured via a lactate dehydrogenase (LDH) release assay. Cells (∼70–80% confluency) were treated with DB33 or DB10 for 24 h. LDH activity was quantified using a formazan-based reaction, with absorbance measured at 500 nm. Cytotoxicity was calculated using the formula: % Cytotoxicity = [(DB10/33) – vehicle) / (tergitol – vehicle)] × 100 where tergitol is the 100% lysis control. Data are shown as the mean ± SD of at least three independent experiments. ∗p ≤ 0.05, ∗∗p ≤ 0.01, ∗∗∗∗p ≤ 0.0001 using a one-way ANOVA with Dunnett’s multiple comparison. DB33-Y was added to *S. aureus*–infected **(B)** RAW264.7 macrophages, **(C)** HMEC-1 endothelial cells or **(D)** bone-marrow-derived macrophages (BMDCs) at 1.5 h post-infection (hpi). Cells were lysed at 12 hpi in (B) and (D) and 8hpi in (C), and intracellular CFUs were quantified to assess the ability of DB33-Y to inhibit bacterial replication. Data represent the mean ± SD of three independent experiments. ∗∗p ≤ 0.01, ****p ≤ 0.0001 by one-way ANOVA with Dunnett’s multiple comparisons.

Given DB33-Y showed reduced cytotoxicity compared to DB10-Y, we selected it for further *in cellulo* testing. To evaluate whether DB33-Y could eradicate intracellular *S. aureus*, we infected RAW264.7 macrophages, bone marrow–derived macrophages (BMDMs), and human microvascular endothelial cells (HMEC-1) and treated cells with DB33-Y. Macrophages and epithelial cells represent key intracellular niches that may be occupied by *S. aureus* during systemic and/or localized infections^50–52^. Here, gentamicin protection assays were performed as previously described^60^. After gentamicin treatment, infected cells were either exposed to vehicle control or DB33-Y at either 100 or 200 μM for approximately 12 h at which the bacterial burden was determined. At 100 μM, DB33-Y reduced intracellular *S. aureus* levels in all three cell types compared to vehicle control (**Figure 7B, C and D**). At 200 μM, intracellular killing was slightly more effective, however, given that this concentration is slightly cytotoxic (**Figure S16A**), the killing of cells is likely influencing bacterial counts. Despite this, these results indicate that DB33-Y is effective at clearing intracellular bacteria at concentrations not toxic to host cells.

### DB33-Y reduces lesion size and bacterial burden in a murine skin infection model

Next, we sought to assess whether DB33-Y would display activity against *S. aureus in vivo*. To this end we employed a murine skin infection model where *S. aureus* can survive and replicate and cause dermonecrotic lesions. Using this model we evaluated the impact of DB33-Y treatment on lesion size and *S. aureus* tissue burden as compared to vehicle control treated animals. Mice were co-injected subcutaneously with USA300 LAC and DB33-Y (100 μM final concentration) or vehicle, followed by a subsequent injection at 24 hours post infection (hpi). At 72 hpi, lesion sizes were significantly smaller in the DB33-Y treated group as compared to control treated animals (**Figure 8A, B**). While bacterial burden was also significantly reduced (**Figure 8C**), the decrease in CFU was modest relative to the reduction in lesion size. This disparity prompted us to explore whether DB33-Y might function, at least in part, through attenuation of virulence rather than by complete bacterial clearance alone.

**Figure 8.**
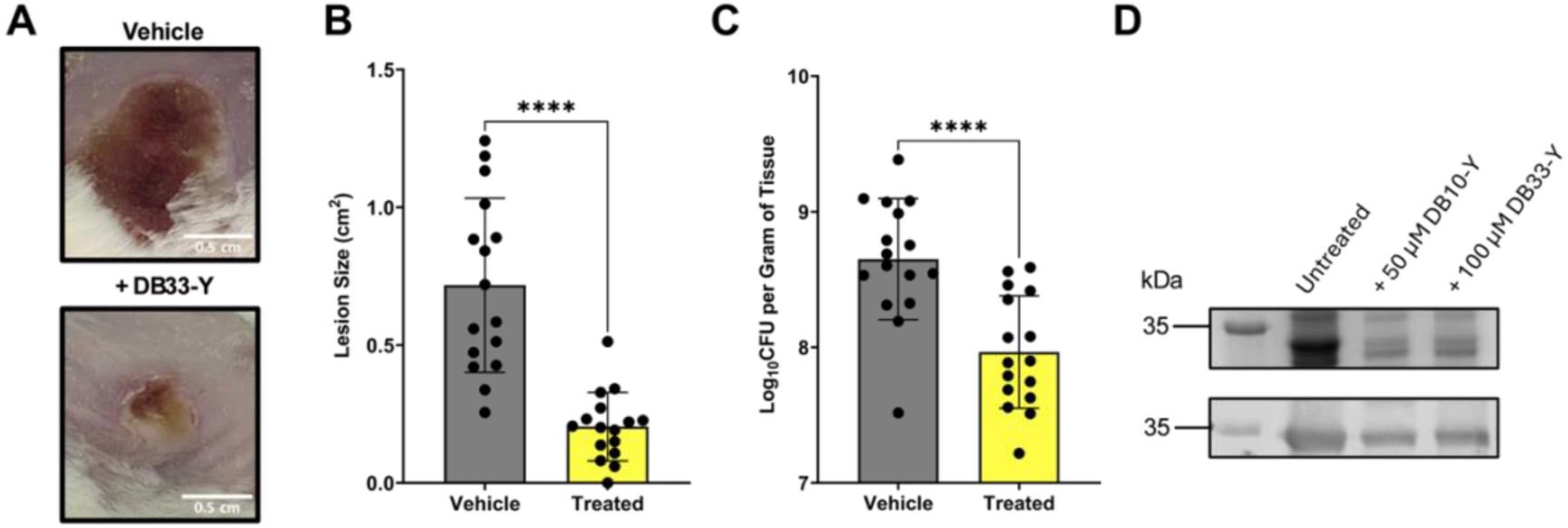
DB33-Y treatment reduces bacterial burden and lesion size in a skin infection model. (**A**) Representative images of skin lesions at 3 days post-infection (dpi). Mice were subcutaneously co-injected with *S. aureus* USA300 LAC (∼5 x 10^7^ CFU) and 100 μM DB33-Y, followed by a second DB33-Y dose (100 μM) at 1 dpi. Mice were sacrificed at 3 dpi and lesion burden was evaluated by quantifying (**B**) CFU/mg of homogenized tissue and **(C**) lesion area. Data represent the mean ± SD of three independent experiments. ∗∗p ≤ 0.01, ∗∗∗∗p ≤ 0.0001 by one-way ANOVA with Dunnett’s multiple comparisons. (**D**) Coomassie-stained gel (top) and Western blot (bottom) showing Hla protein levels following treatment with DB10-Y and DB33-Y at their respective MICs (50 μM and 100 μM).

The *hla* gene encodes α-hemolysin, a pore-forming cytotoxin that contributes to host cell lysis and tissue destruction in *S. aureus* skin infections^61^. Supporting this anti-virulence hypothesis, analysis of secreted proteins by both Coomassie staining and Western blot revealed reduced α-hemolysin levels in both DB10-Y and DB33-Y treated bacteria (**Figure 8D**). This mirrors findings from our previous RNA-seq data with DB10-Y, where *hla* expression was also downregulated. Notably, the full SDS-PAGE gel (**Figure S16B**) shows that additional secreted proteins are also differentially expressed in response to DB10-Y and DB33-Y, indicating broader effects on the bacterial secretome. Collectively, these results suggest that DB33-Y may mitigate tissue damage by suppressing key virulence determinants such as α-hemolysin, alongside other secreted factors.

## DISCUSSION

*Staphylococcus aureus* is a major human pathogen responsible for a wide range of diseases, including skin and soft tissue infections, pneumonia, and bacteremia^1,62^. The rise of methicillin-resistant *S. aureus* (MRSA) has significantly complicated treatment, as these strains are resistant not only to β-lactam antibiotics but also to multiple other drug classes^63^. This growing resistance has underscored the urgent need for novel antimicrobial agents, particularly those that act through mechanisms distinct from conventional antibiotics.

In looking at hits from a previously conducted high-throughput chemical screen^27,28^, we noted that approximately 11% of compounds active against MRSA contained planar, aromatic moieties—structural features commonly associated with DNA intercalation^9^. While such structures are often deprioritized due to concerns over mammalian genotoxicity^21,22^, several, including doxorubicin, have demonstrated potent antimicrobial activity^8,63^. Among the few reproducibly active hits, we prioritized DB10, which contains a fluorene core, for its potent antibacterial activity and unique photochemical properties.

Notably, we observed that upon prolonged ambient light exposure DB10 underwent a spontaneous color change from red (DB10-R) to yellow (DB10-Y), a process we confirmed to be driven specifically by UVA (365 nm) irradiation. UV-Vis spectra revealed a sharp isosbestic point indicating a clean, two-species conversion between DB10-R and DB10-Y, rather than nonspecific photodegradation. Moreover, NMR analysis confirmed that this is a one-way structural rearrangement rather than a reversible isomerization. This irreversible transformation distinguishes DB10 from classical photoswitchable antibiotics, which typically rely on reversible cis–trans isomerization of functional groups such as azobenzenes or stilbenes^64,65^. Furthermore, unlike conventional photoswitches which often necessitate fluorophore conjugation with traditional antibiotics or rely on photodynamic ROS generation^66,67^, DB10 exhibited antibacterial activity independently of these features. Importantly, both DB10-R and DB10-Y possess unique chemical structures that, to our knowledge, have not been previously characterized in the literature.

Photoconversion of DB10-R to DB10-Y involves the loss of the hydrophobic *N*-methylaniline group and formation of a polar aldehyde, resulting in enhanced aqueous solubility and slightly reduced cytotoxicity against RAW264.7 macrophages. Notably, this structural shift is also accompanied by a slight reduction in the MIC across most Gram-positive species tested. While both forms are ineffective against Gram-negative bacteria—likely due to outer membrane impermeability, a common barrier to DNA intercalating agents^6^—they show broad activity against Gram-positive pathogens. However, certain species, including *S. cohnii*, *S. warneri*, and *B. subtilis*, remained less susceptible. This reduced efficacy may reflect species-specific differences in membrane permeability or efflux, but it is also possible that DB10 compounds exhibit sequence-specific DNA binding, and that the relevant target motifs are less prevalent or accessible in these bacterial genomes, as has been observed with other intercalators^8^.

To further elucidate the functional differences following DB10 photoconversion, we performed RNA-seq analysis on USA300 LAC treated with either DB10-R or DB10-Y. Both treatments reduced expression of key virulence factors, including *lukD* (a leukotoxin that lyses immune cells)^68^, *hla* (an alpha-hemolysin that damages host membranes)^69^, and the *spl* operon (serine proteases involved in immune evasion and tissue dissemination)^70^. Western blots confirmed reduced Hla protein levels after DB10-Y or DB33-Y treatment. Repression of these virulence genes after antibiotic exposure has been well-established and may reflect a general stress response or disruption of global regulators^28,68,69,71^.

DB10-R induced upregulation of *clpB*, *clpC*, *mcsA*, and *mcsB*—components of the Clp proteostasis network that aid in refolding and degradation of misfolded proteins during stress^30,72^. Also upregulated was *umuC*, an error-prone polymerase typically activated during DNA damage^73^. Notably, canonical SOS regulators *recA* and *lexA*^7^ were not induced. In contrast to antibiotics like mitomycin C and ciprofloxacin, which activate the SOS cascade via RecA-dependent LexA cleavage^73,74^, DB10-R seems to induce DNA damage in a manner that bypasses classical SOS sensing, possibly involving alternative regulatory pathways or post-transcriptional mechanisms.

DB10-Y elicited a similar, but distinct, RecA-independent stress response. While *clpB* and *clpC* were again upregulated, though slightly below statistical cut-off, DB10-Y–treated cells showed increased expression of *rusA* and *ssb*, which are genes involved in recombinational DNA repair. *rusA* encodes a Holliday junction resolvase capable of resolving recombination intermediates^33,73^, while *ssb* stabilizes single-stranded DNA at stalled replication forks^34^. These genes are not part of the RecA-dependent SOS response, suggesting that DB10-Y promotes replication stress or recombination intermediates that are resolved via RecA-independent mechanisms. One such RecA-independent mechanism, single-strand annealing (SSA), repairs double-strand breaks (DSBs) by aligning and rejoining homologous sequences without requiring strand invasion by RecA^31,75^. While SSA is well characterized in *E. coli* and eukaryotes, its existence and mechanistic details in *S. aureus* remain largely uncharacterized^31^. Given the functions of *rusA* and *ssb*, it is plausible that *S. aureus* employs an SSA-like repair route in response to DB10-Y–induced DNA damage. Moreover, we found that DB10-Y displaced EtBr from DNA, suggesting an intercalative mode of action, and caused double-strand breaks without detectable single-strand breaks. Planar aromatic compounds, like DB10-Y, disrupt DNA by interfering with replication and transcription^76^, often causing DSBs through replication fork collapse or mechanical stress^76,77^. This aligns with increased expression of SSA-related repair genes following DB10-Y exposure.

Given the atypical activation of the DNA damage response observed with DB10-Y, we hypothesized that its mechanism of action is pleiotropic and not limited to classical RecA-dependent DNA damage pathways. To further investigate this, we attempted to evolve resistance through serial passaging. However, only low-level resistance emerged, and only after a prolonged period of selection, suggesting that high-level resistance may be difficult to achieve, likely due to the essential nature of the DNA target.

Interestingly, whole-genome sequencing of resistant populations revealed mutations in *clpX* or the *clpX* upstream promoter region in two independent subpopulations. While *clpX* is not likely the primary target of DB10-Y, its repeated mutation suggests a compensatory role in mitigating the cellular effects of the compound, possibly through modulation of proteostasis or stress responses. The absence of mutations in typical resistance pathways such as efflux pumps, DNA repair genes, or alterations in membrane permeability, commonly observed with antibiotics that cause genotoxic stress and DNA damage^78,79^, supports the idea that DB10-Y activates a non-traditional, RecA-independent DNA damage response, possibly engaging alternative or poorly characterized stress responses in *S. aureus*.

ClpX inactivation has previously been associated with increased resistance to β-lactams and DNA-damaging agents^43,80^. However, the divergent phenotypes of YR-1 and YR-2, one showing increased ampicillin resistance and the other increased susceptibility to vancomycin and ofloxacin, argue against a full loss of ClpX function. The lack of broad cross-resistance, combined with unaltered DB10-Y sensitivity in *clpP* mutants suggests resistance arose from altered regulation of *clpX*, rather than complete inactivation. Supporting this, overexpression of the mutant *clpX* alleles reversed the altered susceptibility phenotypes observed, likely by disrupting proteostasis or interfering with stress adaptation networks.

To investigate structure–activity relationships, several DB10 analogs were tested for DNA binding and antibacterial activity. DB10-Y, DB33-Y, DB42, and DB35 all contain a diene side chain conjugated to the fluorene core, which likely enhances planarity and facilitates DNA intercalation via π–π stacking^81,82^. Indeed, all analogs except DB28 were capable of some DNA intercalation, though DB91 and DB55 were weak. However, only DB33 exhibited antibacterial activity, indicating that intercalation alone is not sufficient for function. Indeed, DB42 and DB35 contain nitro groups that increase polarity and may hinder membrane diffusion^83^, limiting intracellular access despite good DNA affinity. In contrast, DB10 and DB33 lack strongly polar groups, likely facilitating passive uptake^84^. DB55 and DB91 lacked conjugated side chains and instead contained bulky groups (e.g., Br in DB55), which may disrupt planarity and reduce DNA binding. Similarly, DB38 lacks conjugation and electron-donating or -withdrawing groups, precluding effective coordination with DNA.

A closer comparison of DB33-R and DB10-R, which differ only by a terminal phenyl group, shows DB33-R has a slightly lower MIC, suggesting possible steric hindrance of DNA interaction. Interestingly, DB33-Y and DB10-Y both generate the same photoproduct (Figure 6C, compound 2), predicted to be the active species, yet DB33-Y had a higher MIC. This may be due to additional breakdown products from DB33-Y that compete with compound 2 for DNA binding or interfere with its activity.

Despite the reduced potency, DB33-Y exhibited relatively low cytotoxicity compared to DB10-Y against RAW264.7 macrophages, making it a more suitable candidate for *in vivo* evaluation. We therefore selected DB33-Y for further testing in a murine skin infection model. Interestingly, after treatment with DB33-Y we observed only a modest decrease in bacterial CFU, yet a significant reduction in lesion size. Although DB33-Y was administered at its *in vitro* MIC (100 μM) based on testing in MHB, this concentration may not reflect the effective bactericidal dose within skin tissue, which is influenced by host absorption, immune responses, and compound distribution^85^. This suggests DB33-Y may function primarily as an anti-virulence agent at sub-MIC levels, which is further supported by our RNA-seq data that showed downregulation of virulence genes and reduced α-hemolysin levels following DB10-Y. Similar concentration-dependent anti-virulence effects have been reported for clindamycin and tigecycline^86,87^. This anti-virulence activity observed for DB10-Y and DB33-Y may result in clinical advantages by reducing tissue damage and immune activation while lowering the selective pressure for resistance^88^.

Overall, DB10 and DB33 represent promising scaffolds for further antibiotic development. Modest structural changes were sufficient to modulate cytotoxicity and preserve DNA intercalation. Additionally, their dual bactericidal and anti-virulence properties may enable targeted therapeutic strategies depending on infection context. Collectively, the combination of *in vitro* potency against MRSA, *in vivo* efficacy in a skin infection model, and limited resistance development supports further exploration of this scaffold as a potent treatment option for MRSA.

## EXPERIMENTAL PROCEDURES

### Bacterial strains and plasmids

Bacterial strains and plasmids are listed in Table S1. All bacteria were cultured at 37°C with shaking at 200 rpm, with the exception of Streptococcal species, which were grown without shaking. Most Gram-positive bacteria were grown in either tryptic soy broth (TSB) or Mueller-Hinton broth (MHB), unless otherwise stated. Streptococcal species were grown in Todd Hewitt Broth supplemented with 1% yeast extract (THY). Gram-negative species were grown in Luria-Bertani (LB) broth.

### High-throughput screen

The initial high-throughput screen used to identify bioactives of interest was conducted previously^27,28^, and these results were revisited to search for alternative moieties of interest. Of the 965 initial hits, 107 contained a planar aromatic moiety, as identified using a manual search. Planar aromatic moieties identified contained benzothiophene-, indole-, naphthalene, fluorene-, chromene-, phenthrene- and anthrone-based core moieties, all of which have been shown to be involved in DNA damage or intercalation^89–93^. However, in a secondary screen in CDM-G, only three of the initial planar-structure hits retained activity These inhibitors contained a fluorene, indole or phenanthrene core moiety. The indole-containing inhibitor has been previously established to be a ligand that binds to estrogen receptor β and thus was screened out^94^. Given the prevalence of fluorene-containing inhibitors in the initial screen (∼10%) compared to the prevalence of phenanthrene-containing inhibitors (4%), along with the lower MIC associated with the fluorene compound in the secondary screen, we chose to proceed with this compound, which we named “DB10”.

### Determination of Minimum Inhibitory Concentration (MIC)

Serial dilutions of DB10 and DB33 were prepared in Mueller-Hinton broth (MHB) for Gram-positive isolates—except for Streptococcus species, which were grown in Todd-Hewitt broth supplemented with yeast extract (THY)—and in Luria-Bertani (LB) broth for Gram-negative isolates. Bacteria were then added to achieve a starting OD₆₀₀ equivalent of 0.01. Cultures were grown overnight at 37°C with or without shaking as required. The concentration that completely inhibited growth, as measured spectrophotometrically, was determined to be the MIC.

### Bactericidal effect of DB10

An overnight culture of USA300 LAC was washed and inoculated, at a starting OD_600_ equivalent of 0.1, into PBS containing various concentrations of inhibitor. Cultures were incubated overnight at 37°C without shaking before being serially diluted and drop plated onto MHA. After overnight incubation of the plates at 37°C, CFU/mL were enumerated.

### UV-based photoconversion of DB10 and DB33

DB10 or DB33 were stored as a powder away from light at -20°C until use. Just before UV exposure, the compounds were reconstituted in 100% DMSO and subsequently diluted 3:100 in DMSO in a 96-well plate. A spectral scan ranging from 290 nm to 610 nm was conducted before UV exposure, immediately after dilution and at every 5 minutes after until complete conversion was observed (∼20 minutes). UV exposure was conducted by placing the 96-well plate on top of a handheld UVGL-25 compact UV lamp with either UVA (356 nm) or UVB (254 nm) wavelengths switched on. The absorbance readings were corrected for background using DMSO.

### Chemical characterization of the red and yellow forms of DB10 and DB33

NMR spectra for characterization were obtained in DMSO-d6 using a 600 MHz Bruker Ascend 600 instrument. NMR chemical shifts are reported in ppm and are calibrated against the residual solvent signals of DMSO-d6 (2.5 ppm). Coupling constants (J) are given in Hz. Electrospray ionization mass spectrometry (ESI-MS) was performed using a Bruker microOTOF 11 spectrometer. High-performance liquid chromatography (HPLC) was performed using a Waters alliance 2695 separation module connected to a Jupiter 5 µm C18 300 Å LC Column, with a Waters 2487 dual λ absorbance detector detecting at 330 and 350 nm. The samples were injected at 4.0 mg/mL using 25:75 acetonitrile: water to 100% water as eluent over 30 minutes.

Effect of irradiation on 10 g/L DMSO solution of DB10 examined by NMR spectroscopy: DB10 (10mg) was dissolved initially in CDCl_3_, however minimal colour change was observed even after 4 days of exposure to ambient light at 21 °C. Given that the rate for the color change should correlate to the solvent condition and so we tried more polar solvent mixture of DMSO-*d_6_*/D_2_O (6/1, v/v). While the photoconversion was still slow, it was improved relative to CDCl_3_. ^1^H NMR spectra were recorded at various time intervals during ambient light exposure in both CDCl_3_ and DMSO-*d_6_*/D_2_O.

Effect of irradiation on 9 g/L DMSO solution of DB33/1 examined by NMR spectroscopy: DB33 (4.5 mg) was dissolved in 0.5 mL of DMSO-d6. The irradiation with UV light (365 nm) was performed using a 10 W LED light source with 1 A constant current, 400 Hz pulse for 15 hours. Next, the sample was irradiated using a 450 nm blue light for 7.8 hours (36 W LED), followed by 8.2 hours of further irradiation while bubbling air into the sample via a needle. The solution was incubated at 21 °C and 1H NMR spectra were recorded at various time intervals over the cumulative 31 hours of irradiation. The sample was then separated using HPLC. The three main fractions were collected and analyzed by mass spectroscopy.

Characterization of compound 1 (DB33): 1H NMR (600 MHz, DMSO) δ 8.04 (d, J = 7.5 Hz, 1H), 7.88 (d, J = 7.3 Hz, 1H), 7.82 (d, J = 7.3 Hz, 1H), 7.78 (d, J = 7.5 Hz, 1H), 7.41 (d, J = 12.3 Hz, 1H), 7.37 – 7.20 (m, 4H), 6.97 (t, J = 12.0 Hz, 1H), 6.92 – 6.82 (m, 2H), 5.45 (t, J = 12.0 Hz, 1H), 2.89 (s, 6H). 13C NMR (151 MHz, DMSO) δ 149.6, 146.0, 140.3, 138.8, 137.6, 136.8, 132.2, 126.9, 126.7, 126.0, 125.7, 125.0, 124.2, 120.2, 120.0, 119.5, 116.7, 100.0, 40.5. HRMS: calcd [M]+ (C20H19N): 273.1517 Found (ESI): 273.1510

### RNA sequencing

USA300 LAC was sub-cultured into TSB to a final OD_600_ of 0.1 and grown until early exponential phase (OD600 ∼ 1.5). Either vehicle control or sub-MIC DB10-R (50 μM in TSB) or DB10-Y (50 μM in TSB) was then added and the cultures were grown for an additional hour. Cells at an OD_600_ equivalent of 3 were combined with an equal volume of RNA-protect Bacterial Reagent (Qiagen) and pelleted before being frozen at -80°C overnight. Frozen pellets were thawed and resuspended in 750 μL of TE buffer (pH 8) and 50 μL of 1 mg/mL lysostaphin and then incubated at 37°C for 1 h. RNA was extracted using 2 mL of TRI Reagent (Sigma) and 500 μL of BCP (Sigma). After mixing, the aqueous layer was added to an equal volume of ice-cold isopropanol, and the samples were then incubated overnight at -20°C. Samples were pelleted at 4°C for 20 minutes at 11 000 × *g*, resuspended in 500 μL of ice-cold 75% ethanol and pelleted again. Pellets were dried and resuspended in 86 μL of nuclease-free ddH_2_O before treatment with Invitrogen Turbo DNase for 1 h, according to the manufacturer’s instructions. RNA was precipitated using a phenol-chloroform extraction. RNA-sequencing and initial data analysis using TPM (Transcripts Per Million) comparisons were conducted by SeqCenter (Pittsburgh, PA). Further analyses were conducted using the Geneious Prime software package.

### DNA intercalation and damage

DNA used for intercalation and damage assays was isolated from USA300 LAC using phenol-chloroform extractions. To determine the DNA intercalative ability of DB10-Y and DB33-Y, an ethidium bromide (EtBr) displacement assay was performed as described previously^15^, with some modifications. Briefly, EtBr was added to PBS in a 96-well plate to a final concentration of 0.6 μM and genomic DNA was added at a final concentration of 20 ng/μL. Samples were briefly incubated at room temperature for 5 minutes to allow the EtBr to intercalate into DNA before various concentrations of DB10-Y and DB33-Y were added. The plate was then incubated at room temperature in the dark for 30 minutes. Fluorescence spectra (530-700 nm) of samples was measured using a Synergy H4 with an excitation of 525 nm, corresponding to the excitation of EtBr. Wells were background corrected using PBS or DB10-Y/DB33-Y with DNA in the absence of EtBr, as appropriate.

A neutral and alkaline/neutral assay was used to visualize double- and single-stranded DNA breaks (DSBs and SSBs respectively) in USA300 LAC gDNA using previously described methods^40^, with modifications. Briefly, USA300 LAC was sub-cultured into fresh TSB containing various concentrations of DB10 or erythromycin. The cultures were incubated overnight at 37°C with shaking before DNA was phenol-chloroform extracted. To visualize DSBs, gDNA was diluted to a concentration of 500 ng/μL in TE buffer. To visualize alkaline unwinding-sensitive sites corresponding to SSBs, 3 µL of 1 M Na_2_HPO_4_ were added to 20 µL of TE buffer containing 500 ng of gDNA to induce unwinding. 9 µL of 0.1 M HCl was then immediately added to the sample, mixed over ice, and allowed to sit for 4 minutes to allow for DNA renaturation. Loading buffer (0.25% bromophenol blue, 60% glycerol) was added to both conditions, and the samples were run on a 0.8% Tris/Borate/EDTA (TBE) gel with SybrSafe for DNA visualization.

### Detection of superoxide formation

Superoxide formation by USA300 LAC cells after DB10-Y exposure was measured as previously described^28^, with modifications. Briefly, DB10-Y or a DMSO vehicle control was added to 100 µL of USA300 at an OD600 of 0.1 in Hank’s Balanced Salt Solution (HBSS). 500 µL of NBT (1 mg/mL) was added to the samples and incubated at 37°C for 60 minutes. The reaction was stopped by adding 100 µL of 0.1 M HCl and the samples were spun at 5000 rpm for 15 minutes. Pellets were resuspended in equal volumes of DMSO and HBSS. The absorbance levels of the solutions were then read at 575 nm to quantify intracellular ROS production.

### Evolution and characterization of resistant mutants

#### Evolutionary adaptation

Overnight cultures of USA300 LAC were sub-cultured 1:100 daily in TSB with 0.5x MIC DB10-Y (50 µM). MICs were taken periodically to evaluate resistance evolution. After 39 days low-level (2-fold increase) resistance was observed. The culture was passaged for an additional 7 days to ensure stability. The resultant culture was plated on agar to isolate individual colonies, and colonies were patched onto either 5% sheep’s blood agar, or standard method caseinate agar^95^ to determine hemolytic and proteolytic capabilities. Two colonies with distinct phenotypes based on the blood and caseinate agar were selected and designated YR-1 and YR-2.

#### Sequencing of resistant mutants

Genomic DNA was isolated from YR-1 and YR-2 using phenol chloroform extraction and sequenced at SeqCenter (Pittsburgh, PA). DNA sequence reads were paired and mapped to the reference USA300_FPR3757 genome (CP000255.1) using the Geneious software package. Variant analysis was completed using FreeBayes.

#### Cloning and overexpression of mutant ClpX

Genomic DNA was isolated from USA300 LAC by phenol-chloroform extraction. Primers containing a *Kpn*I or *Sac*I restriction site were used to amplify *clpX* from either YR-1 or YR-2 (see Table S2 for primers). The amplified product was ligated into pALC2073 that was also digested using *Kpn*I and *Sac*I. The ligated product was transformed into *E. coli* DH5α. Sequence-confirmed recombinant plasmids were isolated from *E. coli* and electroporated into RN4220. Phage lysate was prepared from the resultant RN4220 strain carrying the p*clpX-*YR-1 and YR-2 plasmids using phage 80α and lysates were used to transduce recipient strains YR-1 and YR-2 with recombinant plasmids with selection on chloramphenicol-containing solid media.

#### TCA precipitation of secreted proteins

Cultures of USA300 LAC were sub-cultured into fresh TSB containing DB10-Y or DB33-Y at various concentrations and cultures were grown to stationary phase overnight. Secreted proteins were precipitated using trichloroacetic acid (TCA). Briefly, supernatant from an OD_600_ equivalent of 6 of culture was added to an equal volume of TCA and incubated at 4°C overnight. Proteins were washed twice with ice-cold 70% ethanol and dried at 37°C before being resuspended in 1× Laemmli SDS-PAGE reducing buffer and boiled for 5 minutes at 100°C. Proteins were loaded onto a 12% SDS-PAGE gel and run at 115V for 10 mins, then 150V for 55 mins. Gels were stained with Coomassie for 1 h and then destained overnight before visualization.

#### Western Blot

Western blot analysis of Hla was conducted by transferring the SDS-PAGE-resolved proteins to a nitrocellulose membrane. The membrane was blocked in PBS containing 0.1% (v/v) Tween 20 and 5% (w/v) skim milk powder. Following blocking, the membrane was incubated overnight with a rabbit polyclonal anti-Hla antibody (1:1000; Sigma-Aldrich), then washed three times with PBS-Tween 20. A donkey anti-rabbit IRDye 800 secondary antibody (1:20,000; Rockland) was applied for 1 hour, followed by three PBS-Tween 20 washes. Imaging was performed using an Odyssey CLx system (Li-Cor Biosciences).

#### Lactate Dehydrogenase (LDH) Release

Quantification of cytotoxicity using LDH release was done using the LDH Cytotoxicity Assay Kit (Cat #MAK529) from Sigma. The manufacturer’s instructions were followed, with modifications. Briefly, RAW264.7 macrophages were seeded into a 96-well plate and grown to a confluency of approximately 60% in RPMI + 5% fetal bovine serum (FBS). Media was removed and replaced with fresh media containing various concentrations of either DB10, DB33, a DMSO vehicle control or a tergitol positive control. Cells were incubated at 37°C for 24 h before 150 µL of Reagent was added to each well and the plate was incubated at room temperature for 10 minutes before formazan production corresponding to LDH release was read at 500 nm. Cytotoxicity was calculated using the following equation: Cytotoxicity= (OD_Sample_-OD_Control_)/(OD_Total Lysis_-OD_Control_) × 100 (%). The plate was read before addition of Reagent and these values were used for background correction.

#### Treatment of macrophage and endothelial cell infections

The efficacy of DB33-Y against intracellular *S. aureus* reservoirs was assessed using procedures as previously described^60^. RAW264.7 macrophages were grown in RPMI + 5% fetal bovine serum (FBS), HMEC-1 cells were grown in MCDB 131 supplemented with 10 mM L-Glutamine, 10 ng/mL epidermal growth factor (EGF), 1 µg/mL hydrocortizone and 10% (v/v) FBS and bone marrow-derived macrophages (BMDMs) were grown in RPMI + 10% FBS. For all cell lines cells were seeded into 12-well plates 24 hours prior to infection. Infections were performed in serum-free media until gentamicin treatment, at which serum-containing media was reintroduced. USA300 LAC was used to infect RAW 264.7 macrophages, BMDMs, and HMEC-1 cells at a multiplicity of infection (MOI) of 10. After a 0.5-hour infection period, 100 μg/mL gentamicin was added to wells and incubated for 1 hour to kill all extracellular bacteria, before cells were gently washed to remove gentamicin. DB33-Y was then added to the culture media to evaluate its ability to restrict intracellular bacterial growth. At 12 hours post-infection (hpi) for RAWs and BMDMs and 8 hpi for HMEC-1s, cells were lysed in 0.1% (v/v) Triton X-100 in sterile PBS. Lysates were plated on TSA for CFU enumeration and compared to a control sample lysed at 1.5 hpi (immediately after gentamicin treatment) to assess intracellular bacterial replication. Fold change was calculated by dividing CFUs at 12 or 8 hpi by CFUs at 1.5 hpi.

#### Mice infections

The mouse study (Animal Use Protocol #2021-090) was approved by the University of Western Ontario Animal Care Committee. Eight-to-ten-week-old male and female BALB/c mice were shaved and depilated using Nair 1 day prior to infection. Overnight cultures (grown in TSB) of *S. aureus* USA300 were subcultured at an OD_600_ of 0.1 in TSB and grown to an OD_600_ of 2.0–2.2. Bacterial cells were pelleted and washed in PBS twice. Bacterial cells were then normalized to an OD_600_ equivalent of 3.7 and 25 μL of the bacterial suspension were mixed with either 25 μL of 200 μM B33 (equivalent to a working concentration of approximately ∼ 1x10^9 CFU/mL, and 100 μM, respectively) or 25 μL of PBS with vehicle. This mixture was immediately injected subcutaneously into the rear flanks of each animal. This dosage was chosen based on the *in vitro* MIC of B33 in MHB. At 24 hpi, either 50 μL of 100 μM B33 or 50 μL of PBS with vehicle were injected into the infection sites. Infected mice were monitored daily for 3 days and sacrificed at 72 hpi. Lesions were excised in PBS with 0.1% (v/v) triton X-100, homogenized in a bullet blender tissue homogenizer (2 × 5 min at speed 6) and serially diluted before being plated onto MSA to enumerate CFUs. Lesion sizes were analyzed in ImageJ.

## Data Availability

RNAseq data can be found in the GEO repository under accession number GSE303558. The authors declare that the data supporting the findings of this study are available within the paper and its supporting Supplementary information files.

## ACKNOWLEDGMENTS

We thank E. Brown from McMaster University for providing the list of hits identified in a previous screen against MSSA.

## AUTHOR CONTRIBUTIONS

AG and DEH conceptualized and designed the study. OME performed the high-throughput screening and analysis. CS and ERG performed the NMR and mass spectrometry. MB, VD, and VB each performed various bacteriology based experiments, including MICs. RSF and EP performed the mouse experiments. AG performed all other experiments and analyses not already listed. AG and DEH wrote the manuscript with input from all authors. DEH, OME and EMG obtained funding.

## REFERENCES

1. Tong, S. Y. C., Davis, J. S., Eichenberger, E., Holland, T. L. & Fowler, V. G. Staphylococcus aureus Infections: Epidemiology, Pathophysiology, Clinical Manifestations, and Management. Clin Microbiol Rev 28, 603–661 (2015).

2. Kaur, D. C. & Chate, S. S. Study of Antibiotic Resistance Pattern in Methicillin Resistant Staphylococcus Aureus with Special Reference to Newer Antibiotic. J Glob Infect Dis 7, 78–84 (2015).

3. Chambers, H. F. & DeLeo, F. R. Waves of Resistance: Staphylococcus aureus in the Antibiotic Era. Nature reviews. Microbiology 7, 629 (2009).

4. Lee, A. S. et al. Methicillin-resistant Staphylococcus aureus. Nat Rev Dis Primers 4, 18033 (2018).

5. Silver, L. L. Challenges of Antibacterial Discovery. Clinical Microbiology Reviews 24, 71–109 (2011).

6. Bolhuis, A. & Aldrich-Wright, J. R. DNA as a target for antimicrobials. Bioorganic Chemistry 55, 51–59 (2014).

7. Cheng, K. et al. Staphylococcus aureus SOS response: Activation, impact, and drug targets. mLife 3, 343–366 (2024).

8. Opperman, T. J. et al. DNA Targeting as a Likely Mechanism Underlying the Antibacterial Activity of Synthetic Bis-Indole Antibiotics. Antimicrobial Agents and Chemotherapy 60, 7067–7076 (2016).

9. 9. Mukherjee, A. & Sasikala, W. D. Chapter One - Drug–DNA Intercalation: From Discovery to the Molecular Mechanism. in Advances in Protein Chemistry and Structural Biology (ed. Karabencheva-Christova, T.) vol. 92 1–62 (Academic Press, 2013).

10. Bie, S. et al. Antibiofilm activity of Plumbagin against Staphylococcus aureus. Sci Rep 15, 7948 (2025).

11. Urpo Kinnunen, P. K. Effects of Anti-neoplastic Agents on the Recovery of Bacteria and Yeasts in an Automated Blood Culture System. Scandinavian Journal of Infectious Diseases (2000) doi:10.1080/00365540050164245.

12. Chowdhury, N., Wood, T. L., Martínez-Vázquez, M., García-Contreras, R. & Wood, T. K. DNA-crosslinker cisplatin eradicates bacterial persister cells. Biotechnology and Bioengineering 113, 1984–1992 (2016).

13. Karkare, S. et al. The Naphthoquinone Diospyrin Is an Inhibitor of DNA Gyrase with a Novel Mechanism of Action. The Journal of Biological Chemistry 288, 5149 (2012).

14. Song, R. et al. Naphthoquinone-derivative as a synthetic compound to overcome the antibiotic resistance of methicillin-resistant S. aureus. Commun Biol 3, 1–11 (2020).

15. Chung, B., Kwon, O.-S., Shin, J. & Oh, K.-B. Antibacterial Activity and Mode of Action of Lactoquinomycin A from Streptomyces bacillaris. Marine Drugs 19, 7 (2020).

16. Zhang, H., Lin, J., Rasheed, S. & Zhou, C. Design, synthesis, and biological evaluation of novel benzimidazole derivatives and their interaction with calf thymus DNA and synergistic effects with clinical drugs. Sci. China Chem. 57, 807–822 (2014).

17. Choi, S., Larson, M. A., Hinrichs, S. H. & Narayanasamy, P. Development of potential broad spectrum antimicrobials using *C*2-symmetric 9-fluorenone alkyl amine. Bioorganic & Medicinal Chemistry Letters 26, 1997–1999 (2016).

18. Mishra, N. M. et al. Iterative Chemical Engineering of Vancomycin Leads to Novel Vancomycin Analogs With a High in Vitro Therapeutic Index. Front. Microbiol. 9, (2018).

19. Paul, P. et al. 1,4-Naphthoquinone accumulates reactive oxygen species in Staphylococcus aureus: a promising approach towards effective management of biofilm threat. Arch Microbiol 203, 1183– 1193 (2021).

20. Peng, Q., Tang, X., Dong, W., Sun, N. & Yuan, W. A Review of Biofilm Formation of Staphylococcus aureus and Its Regulation Mechanism. Antibiotics 12, 12 (2023).

21. Buckner, M. M. C., Ciusa, M. L. & Piddock, L. J. V. Strategies to combat antimicrobial resistance: anti-plasmid and plasmid curing. FEMS Microbiology Reviews 42, 781–804 (2018).

22. Suckling, C. From Multiply Active Natural Product To Candidate Drug? Antibacterial (And Other) Minor Groove Binders for DNA. Future Medicinal Chemistry 4, 971–989 (2012).

23. Sakthikumar, K., Isamura, B. K. & Maçedo Krause, R. W. Exploring the antioxidant, antimicrobial, cytotoxic and biothermodynamic properties of novel morpholine derivative bioactive Mn( ii ), Co( ii ) and Ni( ii ) complexes – combined experimental and theoretical measurements towards DNA/BSA/SARS-CoV-2 3CL Pro. RSC Medicinal Chemistry 14, 1667–1697 (2023).

24. Kalepu, S. & Nekkanti, V. Insoluble drug delivery strategies: review of recent advances and business prospects. Acta Pharmaceutica Sinica. B 5, 442 (2015).

25. Crane, J. K., Alvarado, C. L. & Sutton, M. D. Role of the SOS Response in the Generation of Antibiotic Resistance In Vivo. Antimicrobial Agents and Chemotherapy 65, e00013 (2021).

26. Podlesek, Z. & Bertok, D. Ž. The DNA Damage Inducible SOS Response Is a Key Player in the Generation of Bacterial Persister Cells and Population Wide Tolerance. Frontiers in Microbiology 11, 1785 (2020).

27. Bæk, K. T. et al. A Staphylococcus aureus clpX Mutant Used as a Unique Screening Tool to Identify Cell Wall Synthesis Inhibitors that Reverse β-Lactam Resistance in MRSA. Front Mol Biosci 8, 691569 (2021).

28. Gaudreau, A. et al. Mechanistic insights and in vivo efficacy of thiosemicarbazones against methicillin-resistant Staphylococcus aureus. J Biol Chem 107689 (2024) doi:10.1016/j.jbc.2024.107689.

29. Chandrangsu, P., Rensing, C. & Helmann, J. D. Metal Homeostasis and Resistance in Bacteria. Nat Rev Microbiol 15, 338–350 (2017).

30. Frees, D. et al. Clp ATPases are required for stress tolerance, intracellular replication and biofilm formation in Staphylococcus aureus. Molecular Microbiology 54, 1445–1462 (2004).

31. Ha, K. P. & Edwards, A. M. DNA Repair in Staphylococcus aureus. Microbiology and Molecular Biology Reviews : MMBR 85, e00091 (2021).

32. Elsholz, A. K. W., Michalik, S., Zühlke, D., Hecker, M. & Gerth, U. CtsR, the Gram-positive master regulator of protein quality control, feels the heat. The EMBO Journal 29, 3621 (2010).

33. Sharples, G. J., Ingleston, S. M. & Lloyd, R. G. Holliday Junction Processing in Bacteria: Insights from the Evolutionary Conservation of RuvABC, RecG, and RusA. Journal of Bacteriology 181, 5543–5550 (1999).

34. Chen, K.-L., Cheng, J.-H., Lin, C.-Y., Huang, Y.-H. & Huang, C.-Y. Characterization of single-stranded DNA-binding protein SsbB from Staphylococcus aureus: SsbB cannot stimulate PriA helicase. RSC Adv. 8, 28367–28375 (2018).

35. Stahlhut, S. G. et al. The ClpXP protease is dispensable for degradation of unfolded proteins in Staphylococcus aureus. Sci Rep 7, 11739 (2017).

36. Zhou, C. & Fey, P. D. The Acid Response Network of Staphylococcus aureus. Current opinion in microbiology 55, 67 (2020).

37. Davies, M. L. et al. In Depth Analysis of the Quenching of Three Fluorene–Phenylene-Based Cationic Conjugated Polyelectrolytes by DNA and DNA Bases. J. Phys. Chem. B 118, 460–469 (2014).

38. Nishimura, T. et al. DNA Binding of Tilorone: 1H NMR and Calorimetric Studies of the Intercalation. Biochemistry 46, 8156–8163 (2007).

39. Galindo-Murillo, R. & Thomas E Cheatham, I. I. I. Ethidium bromide interactions with DNA: an exploration of a classic DNA–ligand complex with unbiased molecular dynamics simulations. Nucleic Acids Research 49, 3735 (2021).

40. Londero, J. E. L., Schavinski, C. R., Silva, F. D. da, Piccoli, B. C. & Schuch, A. P. Development of a rapid electrophoretic assay for genomic DNA damage quantification. Ecotoxicol Environ Saf 210, 111859 (2021).

41. Dwyer, D. J. et al. Antibiotics induce redox-related physiological alterations as part of their lethality. Proceedings of the National Academy of Sciences 111, E2100–E2109 (2014).

42. Páez, P. L., Becerra, M. C. & Albesa, I. Effect of the association of reduced glutathione and ciprofloxacin on the antimicrobial activity in Staphylococcus aureus. FEMS Microbiology Letters 303, 101–105 (2010).

43. Jensen, C., Fosberg, M. J., Thalsø-Madsen, I., Bæk, K. T. & Frees, D. Staphylococcus aureus ClpX localizes at the division septum and impacts transcription of genes involved in cell division, T7-secretion, and SaPI5-excision. Sci Rep 9, 16456 (2019).

44. Glover, S. A. & Schumacher, R. R. Mutagenicity of *N*-acyloxy-*N*-alkoxyamides as an indicator of DNA intercalation: The role of fluorene and fluorenone substituents as DNA intercalators. Mutation Research/Genetic Toxicology and Environmental Mutagenesis 863–864, 503299 (2021).

45. Fukushima, H. et al. Cyanine Phototruncation Enables Spatiotemporal Cell Labeling. J Am Chem Soc 144, 11075–11080 (2022).

46. Matikonda, S. S. et al. Defining the Basis of Cyanine Phototruncation Enables a New Approach to Single-Molecule Localization Microscopy. ACS Cent. Sci. 7, 1144–1155 (2021).

47. Garavelli, M., Bernardi, F., Olivucci, M. & Robb, M. A. DFT Study of the Reactions between Singlet-Oxygen and a Carotenoid Model. J. Am. Chem. Soc. 120, 10210–10222 (1998).

48. Helmerich, D. A., Beliu, G., Matikonda, S. S., Schnermann, M. J. & Sauer, M. Photoblueing of organic dyes can cause artifacts in super-resolution microscopy. Nat Methods 18, 253–257 (2021).

49. Maoka, T. Chapter 20 - Oxidation products of astaxanthin: An overview. in Global Perspectives on Astaxanthin (eds. Ravishankar, G. A. & Ranga Rao, A.) 411–425 (Academic Press, 2021). doi:10.1016/B978-0-12-823304-7.00013-1.

50. Flannagan, R. S. & Heinrichs, D. E. Macrophage-driven nutrient delivery to phagosomal Staphylococcus aureus supports bacterial growth. EMBO Rep 21, e50348 (2020).

51. Krauss, J. L. et al. Staphylococcus aureus Infects Osteoclasts and Replicates Intracellularly. mBio 10, e02447 (2019).

52. Rollin, G. et al. Intracellular Survival of Staphylococcus aureus in Endothelial Cells: A Matter of Growth or Persistence. Frontiers in Microbiology 8, 1354 (2017).

54. Fauerharmel-Nunes, T. et al. MRSA Isolates from Patients with Persistent Bacteremia Generate Nonstable Small Colony Variants In Vitro within Macrophages and Endothelial Cells during Prolonged Vancomycin Exposure. Infect Immun 91, e0042322 (2023).

54. Flannagan, R. S., Farrell, T. J., Trothen, S. M., Dikeakos, J. D. & Heinrichs, D. E. Rapid removal of phagosomal ferroportin in macrophages contributes to nutritional immunity. Blood Adv 5, 459–474 (2021).

55. Flannagan, R. S., Heit, B. & Heinrichs, D. E. Intracellular replication of Staphylococcus aureus in mature phagolysosomes in macrophages precedes host cell death, and bacterial escape and dissemination. Cell Microbiol 18, 514–535 (2016).

56. Goncheva, M. I., Flannagan, R. S. & Heinrichs, D. E. De Novo Purine Biosynthesis Is Required for Intracellular Growth of Staphylococcus aureus and for the Hypervirulence Phenotype of a purR Mutant. Infect Immun 88, e00104–20 (2020).

57. Flannagan, R. S. & Heinrichs, D. E. A Fluorescence Based-Proliferation Assay for the Identification of Replicating Bacteria Within Host Cells. Front Microbiol 9, 3084 (2018).

58. Flannagan, R. S., Kuiack, R. C., McGavin, M. J. & Heinrichs, D. E. Staphylococcus aureus Uses the GraXRS Regulatory System To Sense and Adapt to the Acidified Phagolysosome in Macrophages. mBio 9, e01143–18 (2018).

59. Kumar, P., Nagarajan, A. & Uchil, P. D. Analysis of Cell Viability by the Lactate Dehydrogenase Assay. Cold Spring Harb Protoc 2018, (2018).

60. El-Halfawy, O. M. et al. Discovery of an antivirulence compound that reverses β-lactam resistance in MRSA. Nat Chem Biol 16, 143–149 (2020).

61. Tam, K. & Torres, V. J. Staphylococcus aureus Secreted Toxins and Extracellular Enzymes. Microbiology Spectrum 7, 10.1128/microbiolspec.gpp3 (2019).

62. Chen, H. et al. Exploring the Role of Staphylococcus aureus in Inflammatory Diseases. Toxins (Basel) 14, 464 (2022).

63. Liu, W.-T. et al. Emerging resistance mechanisms for 4 types of common anti-MRSA antibiotics in Staphylococcus aureus: A comprehensive review. Microb Pathog 156, 104915 (2021).

64. Aubert, A., Fayeulle, A., Vayssade, M., Billamboz, M. & Léonard, E. New Trends on Photoswitchable Antibiotics: From Syntheses to Applications. Photocatalysis: Research and Potential 1, 10007 (2023).

65. Velema, W. A. et al. Optical control of antibacterial activity. Nature Chem 5, 924–928 (2013).

66. Martino, M. D. et al. Azobenzene as Antimicrobial Molecules. Molecules 27, 5643 (2022).

67. Pérez-Aranda, M. et al. Antimicrobial and Antibiofilm Effect of 4,4′-Dihydroxy-azobenzene against Clinically Resistant Staphylococci. Antibiotics 11, 1800 (2022).

68. Yang, H. et al. Anti-staphylococcus Antibiotics Interfere With the Transcription of Leucocidin ED Gene in Staphylococcus aureus Strain Newman. Front. Microbiol. 11, (2020).

69. Hodille, E. et al. The Role of Antibiotics in Modulating Virulence in Staphylococcus aureus. Clinical Microbiology Reviews 30, 887–917 (2017).

70. Pietrocola, G., Nobile, G., Rindi, S. & Speziale, P. Staphylococcus aureus Manipulates Innate Immunity through Own and Host-Expressed Proteases. Frontiers in Cellular and Infection Microbiology 7, 166 (2017).

71. Stevens, D. L. et al. Impact of Antibiotics on Expression of Virulence-Associated Exotoxin Genes in Methicillin-Sensitive and Methicillin-Resistant Staphylococcus aureus. The Journal of Infectious Diseases 195, 202–211 (2007).

72. Wozniak, D. J., Tiwari, K. B., Soufan, R. & Jayaswal, R. K. The mcsB gene of the clpC operon is required for stress tolerance and virulence in Staphylococcus aureus. Microbiology 158, 2568 (2012).

73. Anderson, K. L. et al. Characterization of the Staphylococcus aureus Heat Shock, Cold Shock, Stringent, and SOS Responses and Their Effects on Log-Phase mRNA Turnover. Journal of Bacteriology 188, 6739 (2006).

74. Cirz, R. T. et al. Complete and SOS-Mediated Response of Staphylococcus aureus to the Antibiotic Ciprofloxacin. J Bacteriol 189, 531–539 (2007).

75. Bhargava, R., Onyango, D. O. & Stark, J. M. Regulation of Single Strand Annealing and its role in genome maintenance. Trends in genetics : TIG 32, 566 (2016).

76. El-Adl, K., El-Helby, A.-G. A., Sakr, H. & Elwan, A. Design, synthesis, molecular docking and anti-proliferative evaluations of [1,2,4]triazolo[4,3-*a*]quinoxaline derivatives as DNA intercalators and Topoisomerase II inhibitors. Bioorganic Chemistry 105, 104399 (2020).

77. Saxena, S. & Zou, L. Hallmarks of DNA Replication Stress. Molecular cell 82, 2298 (2022).

78. Patel, D., Kosmidis, C., Seo, S. M. & Kaatz, G. W. Ethidium Bromide MIC Screening for Enhanced Efflux Pump Gene Expression or Efflux Activity in Staphylococcus aureus. Antimicrobial Agents and Chemotherapy 54, 5070 (2010).

79. Suzuki, T. et al. Wall teichoic acid protects Staphylococcus aureus from inhibition by Congo red and other dyes. Journal of Antimicrobial Chemotherapy 67, 2143–2151 (2012).

80. Bæk, K. T. et al. β-Lactam Resistance in Methicillin-Resistant Staphylococcus aureus USA300 Is Increased by Inactivation of the ClpXP Protease. Antimicrobial Agents and Chemotherapy 58, 4593– 4603 (2014).

81. Barone, G. et al. Intercalation of Daunomycin into Stacked DNA Base Pairs. DFT Study of an Anticancer Drug. Journal of Biomolecular Structure and Dynamics 26, 115–129 (2008).

82. Neto, B. A. D. & Lapis, A. A. M. Recent Developments in the Chemistry of Deoxyribonucleic Acid (DNA) Intercalators: Principles, Design, Synthesis, Applications and Trends. Molecules 14, 1725 (2009).

83. Olender, D., Żwawiak, J. & Zaprutko, L. Multidirectional Efficacy of Biologically Active Nitro Compounds Included in Medicines. Pharmaceuticals 11, 54 (2018).

84. Yang, N. J. & Hinner, M. J. Getting Across the Cell Membrane: An Overview for Small Molecules, Peptides, and Proteins. Methods in molecular biology (Clifton, N.J.) 1266, 29 (2015).

85. Smith, R., Russo, J., Fiegel, J. & Brogden, N. Antibiotic Delivery Strategies to Treat Skin Infections When Innate Antimicrobial Defense Fails. Antibiotics 9, 56 (2020).

86. Andersson, D. I. & Hughes, D. Microbiological effects of sublethal levels of antibiotics. Nat Rev Microbiol 12, 465–478 (2014).

87. Shi, J. et al. Sub-inhibitory concentrations of tigecycline could attenuate the virulence of Staphylococcus aureus by inhibiting the product of α-toxin. Microbiology Spectrum 13, e01344 (2025).

88. Rasko, D. A. & Sperandio, V. Anti-virulence strategies to combat bacteria-mediated disease. Nat Rev Drug Discov 9, 117–128 (2010).

89. Aleksić, M. et al. Novel Substituted Benzothiophene and Thienothiophene Carboxanilides and Quinolones: Synthesis, Photochemical Synthesis, DNA-Binding Properties, Antitumor Evaluation and 3D-Derived QSAR Analysis. J. Med. Chem. 55, 5044–5060 (2012).

90. Vincent, S. et al. A Fluorogenic Covalent Chromone-Based Intercalator with a Mega-Stokes Shift for Sensing DNA Hybridization. Chemosensors 11, 161 (2023).

91. Lauria, A., La Monica, G., Bono, A. & Martorana, A. Quinoline anticancer agents active on DNA and DNA-interacting proteins: From classical to emerging therapeutic targets. European Journal of Medicinal Chemistry 220, 113555 (2021).

92. Chen, K. X., Gresh, N. & Pullman, B. A theoretical study of anthracene and phenanthrene derivatives acting as A-T specific intercalators. Nucleic Acids Research 14, 9103 (1986).

93. Wei, Y., Zhang, D., Pan, J., Gong, D. & Zhang, G. Elucidating the Interaction of Indole-3-Propionic Acid and Calf Thymus DNA: Multispectroscopic and Computational Modeling Approaches. Foods 13, 1878 (2024).

94. Chesworth, R. et al. Estrogen receptor β selective ligands: Discovery and SAR of novel heterocyclic ligands. Bioorganic & Medicinal Chemistry Letters 15, 5562–5566 (2005).

95. Martley, F. G., Jayashankar, S. R. & Lawrence, R. C. An Improved Agar Medium for the Detection of Proteolytic Organisms in Total Bacterial Counts. Journal of Applied Bacteriology 33, 363–370 (1970).

