## Supplemental for "Photochemical modification of two fluorene-based molecules yields structurally distinct DNA intercalators with potent anti-MRSA activity"

### Supplementary Information

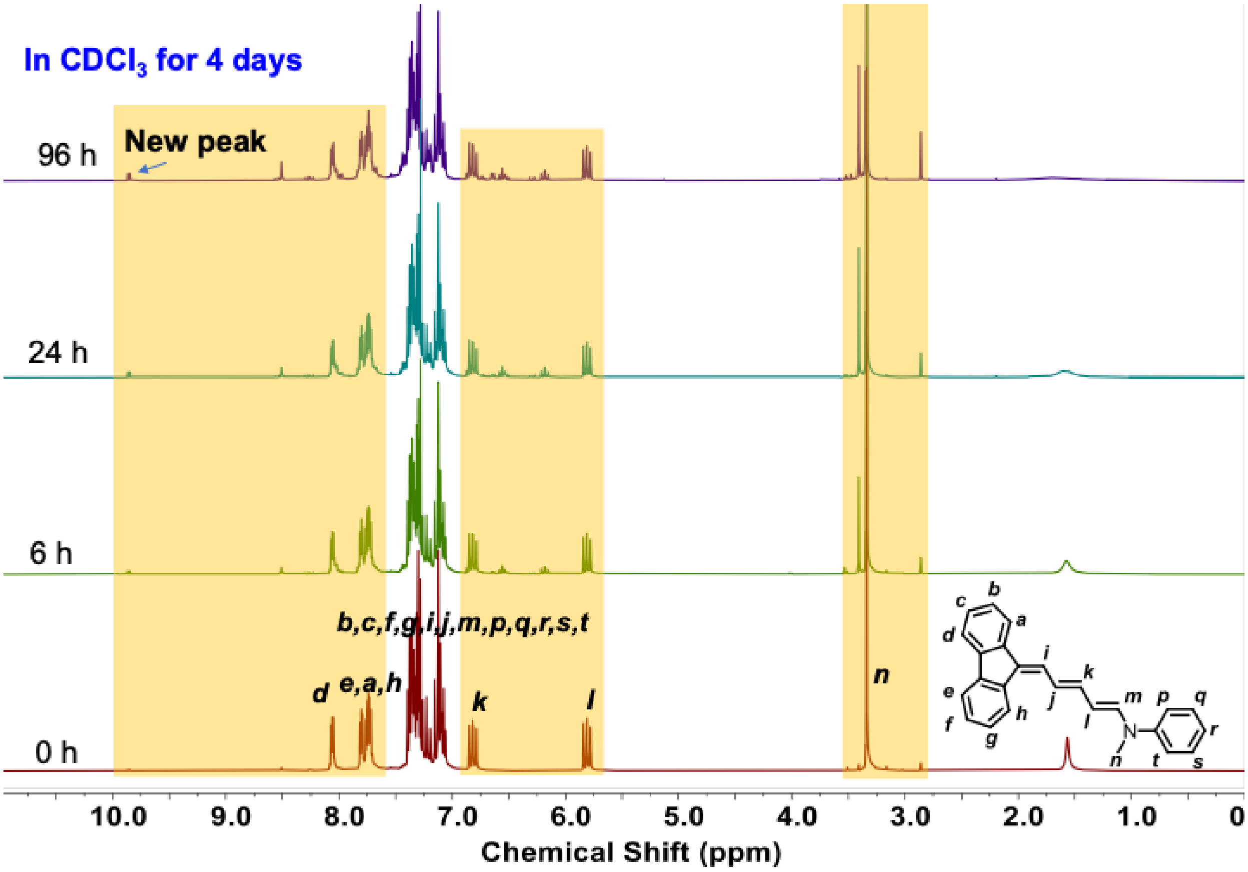


**Figure S1.** $^1$**H NMR spectrum of DB10 incubated at 21ºC over 4 days ($CDCl_3$, 600 MHz).**

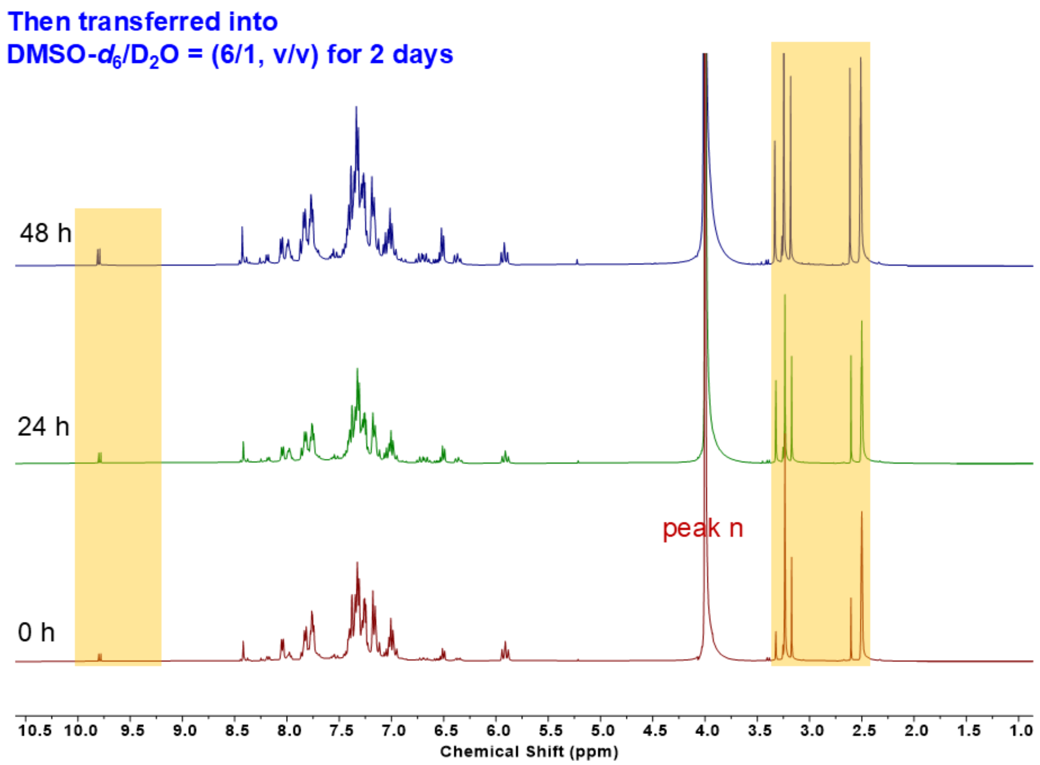


**Figure S2.** $^1$H NMR spectrum of DB10 incubated in 6:1 DMSO-$d_6$: $D_2O$ over 2 days after incubation in $CDCl_3$ for 4 days at 21ºC (600 MHz).

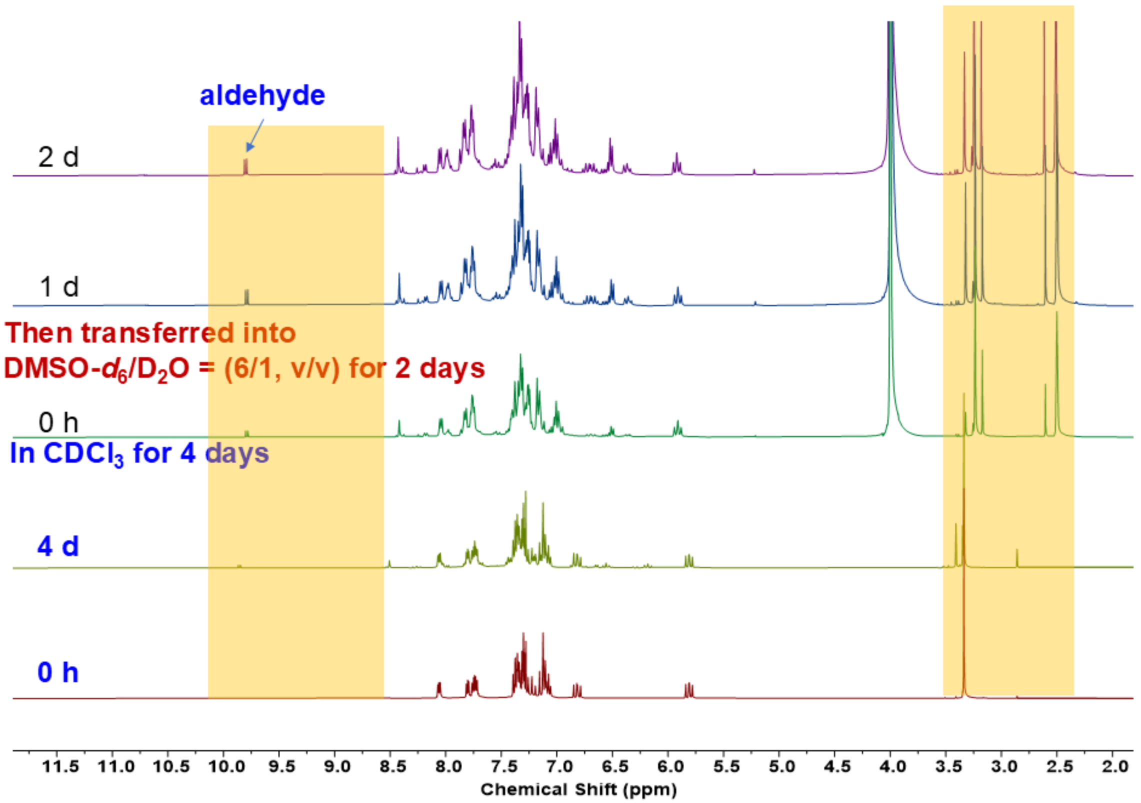

**Figure S3. Overlay of $^1$H NMR spectrum of DB10 incubated in $CDCl_3$ and 6:1 DMSO-$d_6$: $D_2O$ over 6 days at 21°C showing emergence of the aldehyde peak at 9.8 ppm (600 MHz).**

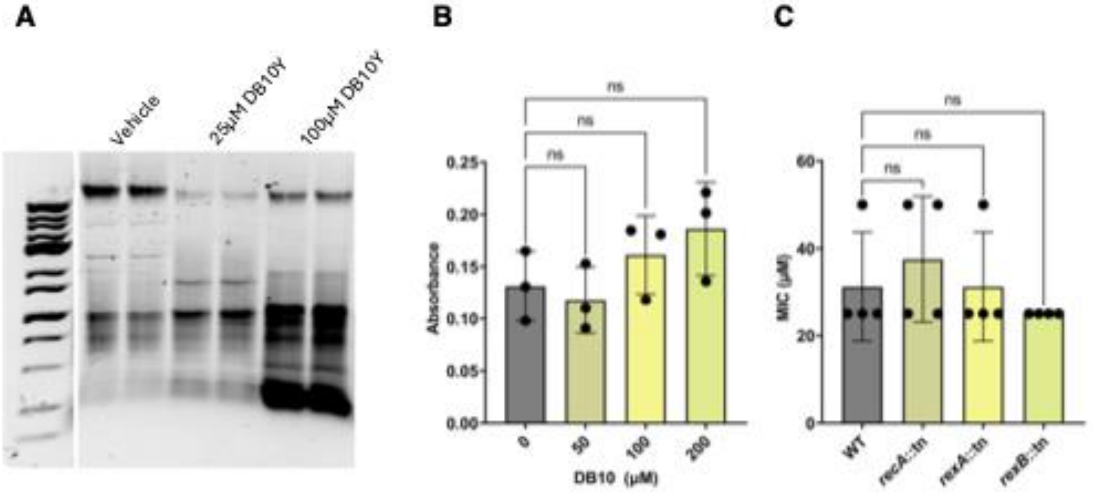


**Figure S4: Exposure of *S. aureus* to DB10-Y results in non-RecA-mediated DNA damage response.** (**A**) Neutral/alkaline treatment of genomic DNA isolated from *S. aureus* cultures grown in the presence of either DMSO vehicle control, or DB10-Y at the indicated concentrations. Duplicate samples were run through a 0.8% Tris/Borate/EDTA (TBE) gel. (**B**) Intracellular superoxide ($O2^{\bullet-}$) levels in USA300 LAC after exposure to DB10-Y for 1h. Nitro-blue tetrazolium (NBT) reduction was used to determine $O2^{\bullet-}$ levels. (**C**) MIC of USA300 LAC compared to mutants with transposon insertions in genes, as indicated, involved in the canonical DNA damage response. Data are shown as the mean ± SD of at least three independent experiments.

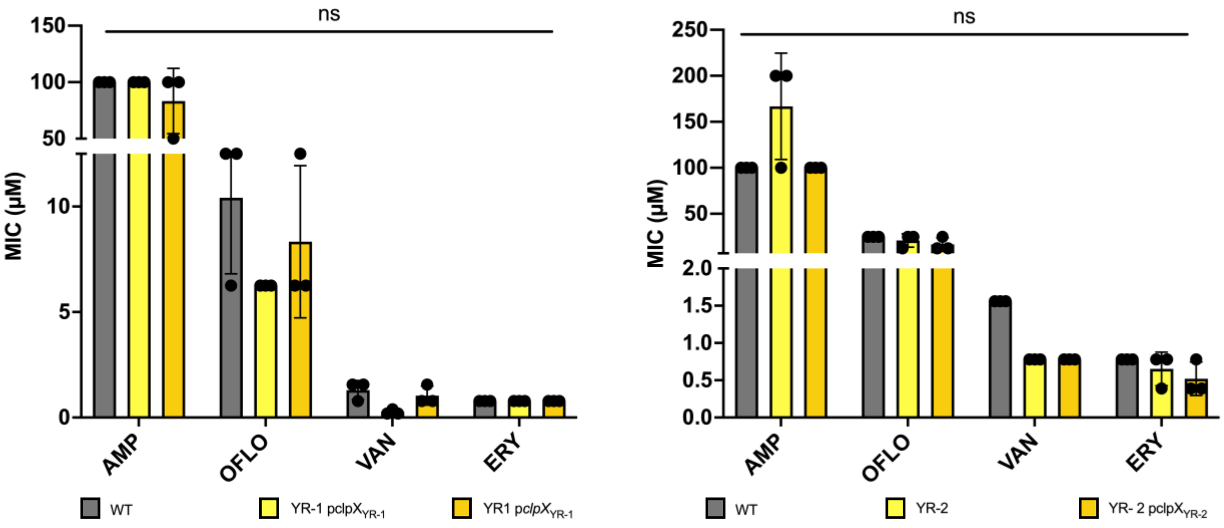


**Figure S5: Overexpression of *clpX* alters bacterial susceptibility to classical antibiotics.** MICs of (**A**) YR-1 and (**B**) YR-2 compared to WT USA300 LAC and their corresponding *clpX* overexpression strains against antibiotics previously associated with *clpX*-linked resistance (AMP) and antibiotics involved in DNA damage (OFLO and VAN) and downstream effects (ERY).

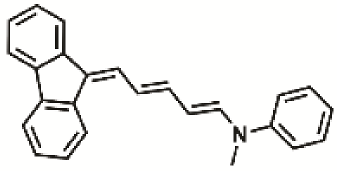

DB10

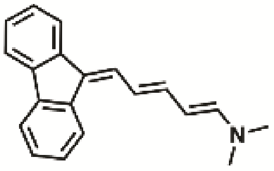

DB33

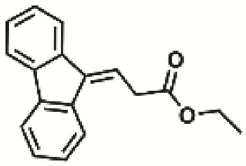

DB91

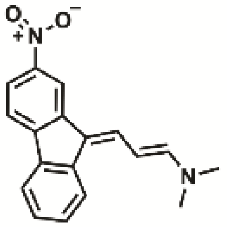


DB35

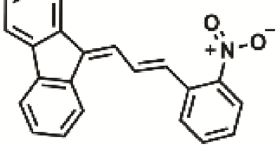


DB42

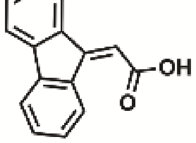


DB28

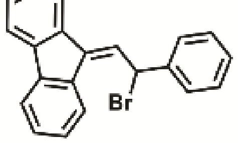


DB55

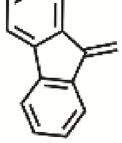

DB38

**Figure S6 Structures of analogs of DB10.** Compounds with biological activity are labelled in red, and compounds without biological activity are labelled in black.

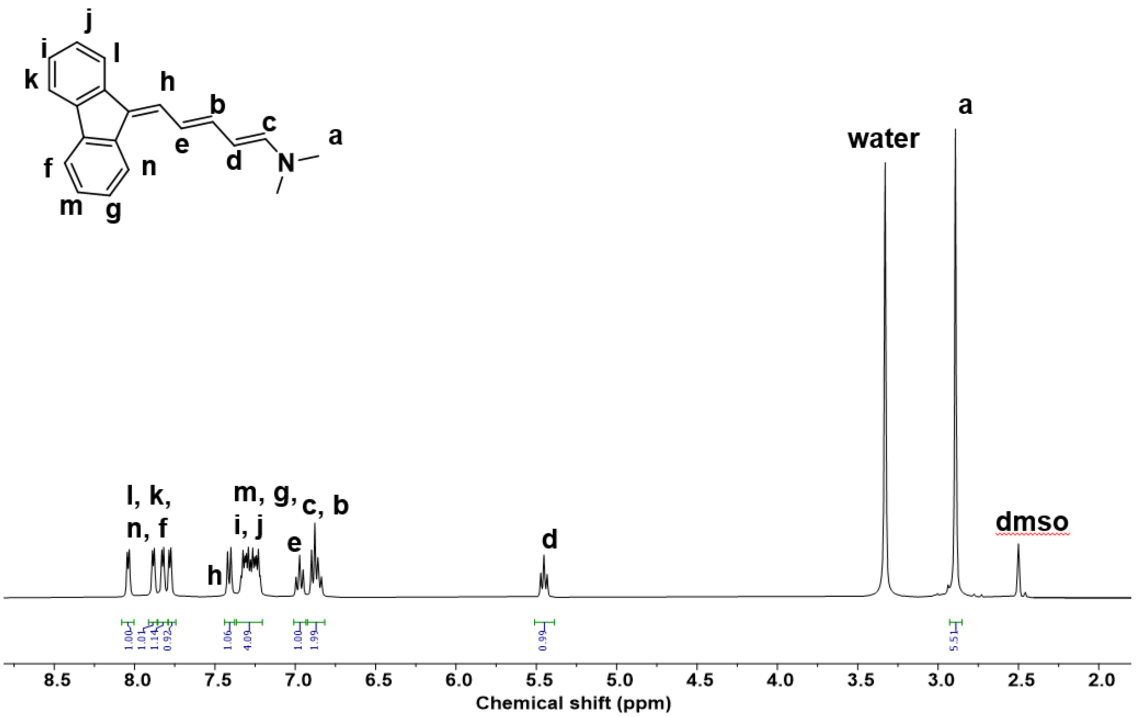


**Figure S7. [1]H NMR spectrum of compound 1 (600 MHz, DMSO-$d_6$)**

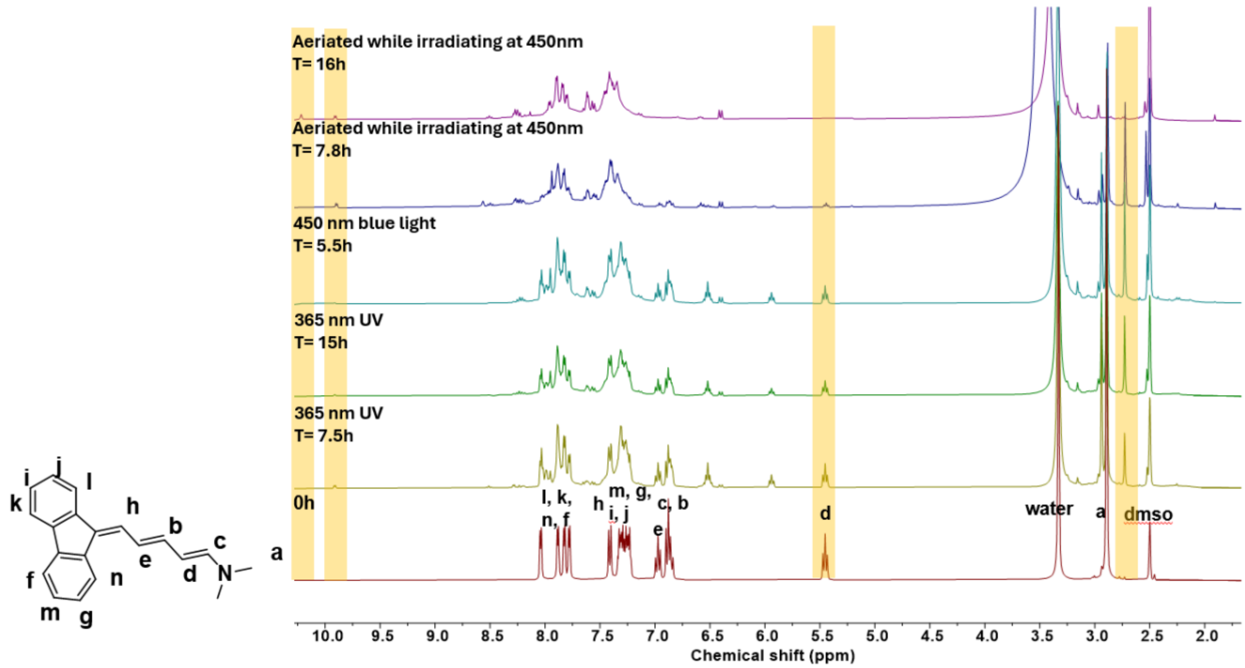

**Figure S8. $^1$H NMR spectra of compound 1 (DB33) over 31 hours of irradiation, incubated at 21 °C (600 MHz, DMSO-$d_6$).**

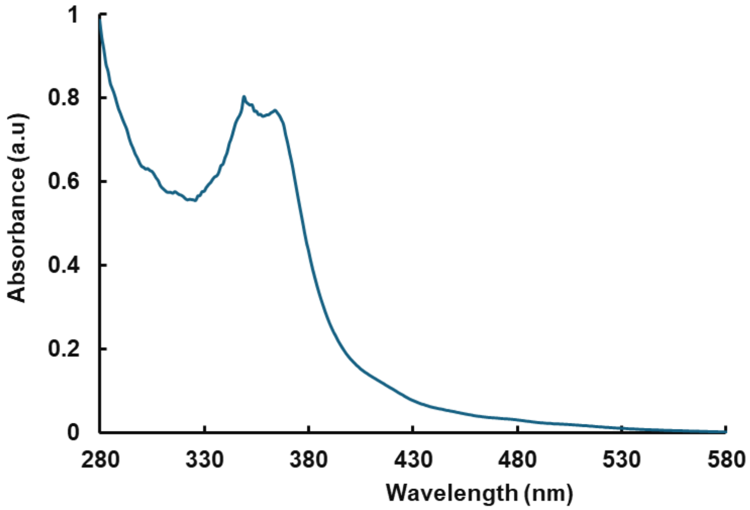


**Figure S9. UV-visible spectrum of the photochemical degradation products of compound 1 (DB33) after a total of 31 hours of irradiation.**

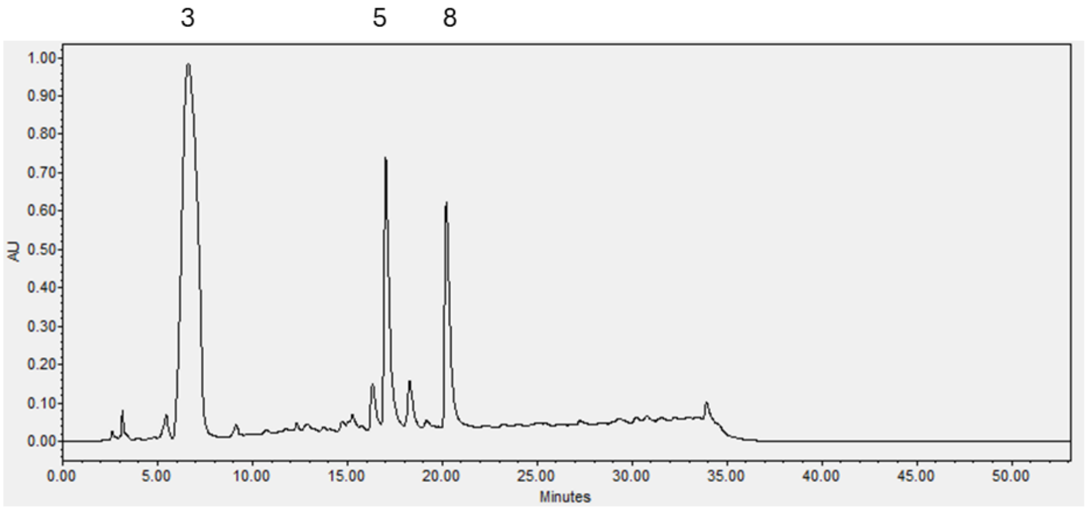


**Figure S10. HPLC trace of the photodegradation products from the final NMR sample in Figure S7.**

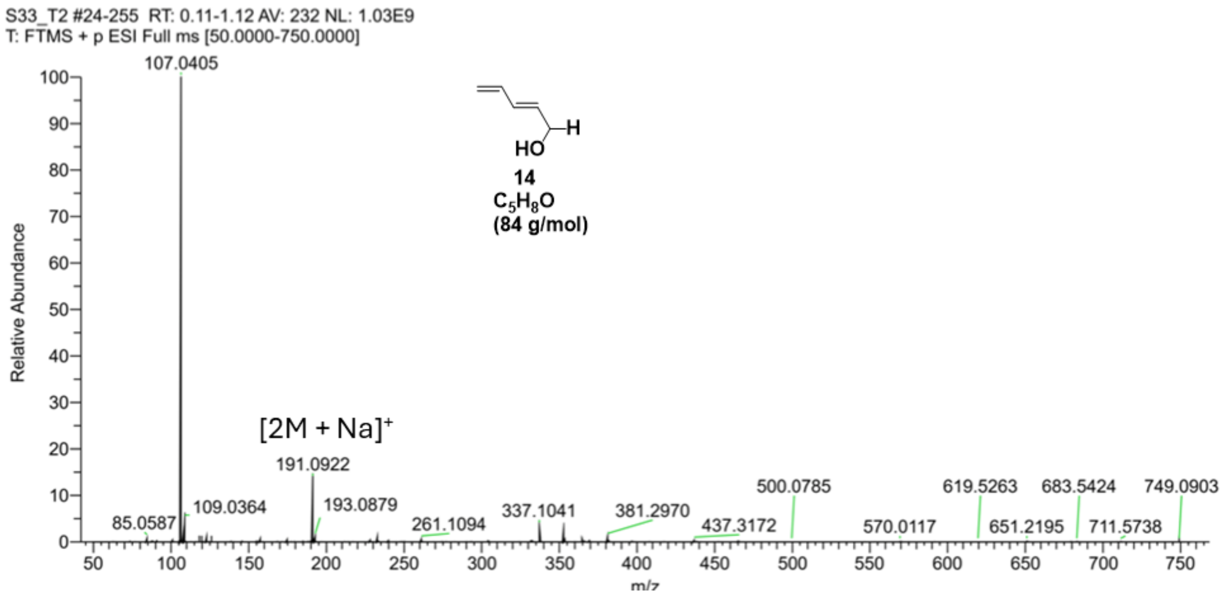


**Figure S11. Mass spectrum of peak 3 from the HPLC separation in Figure S10.**

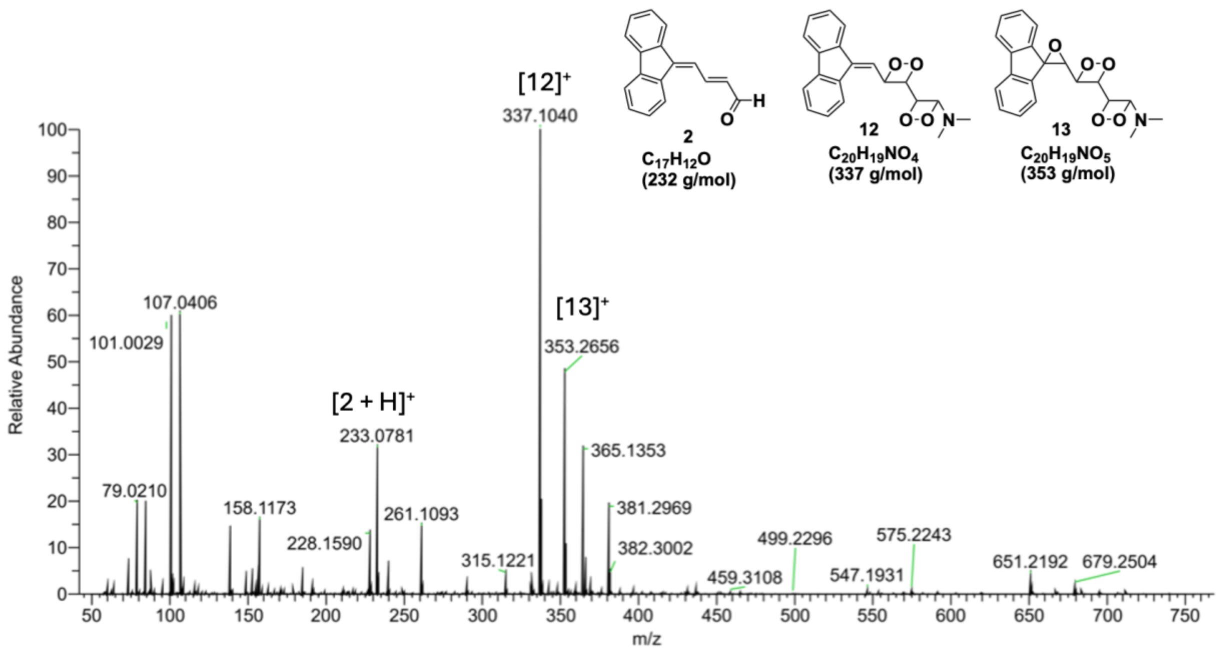


**Figure S12. Mass spectrum of peak 5 from the HPLC separation in Figure S10.**

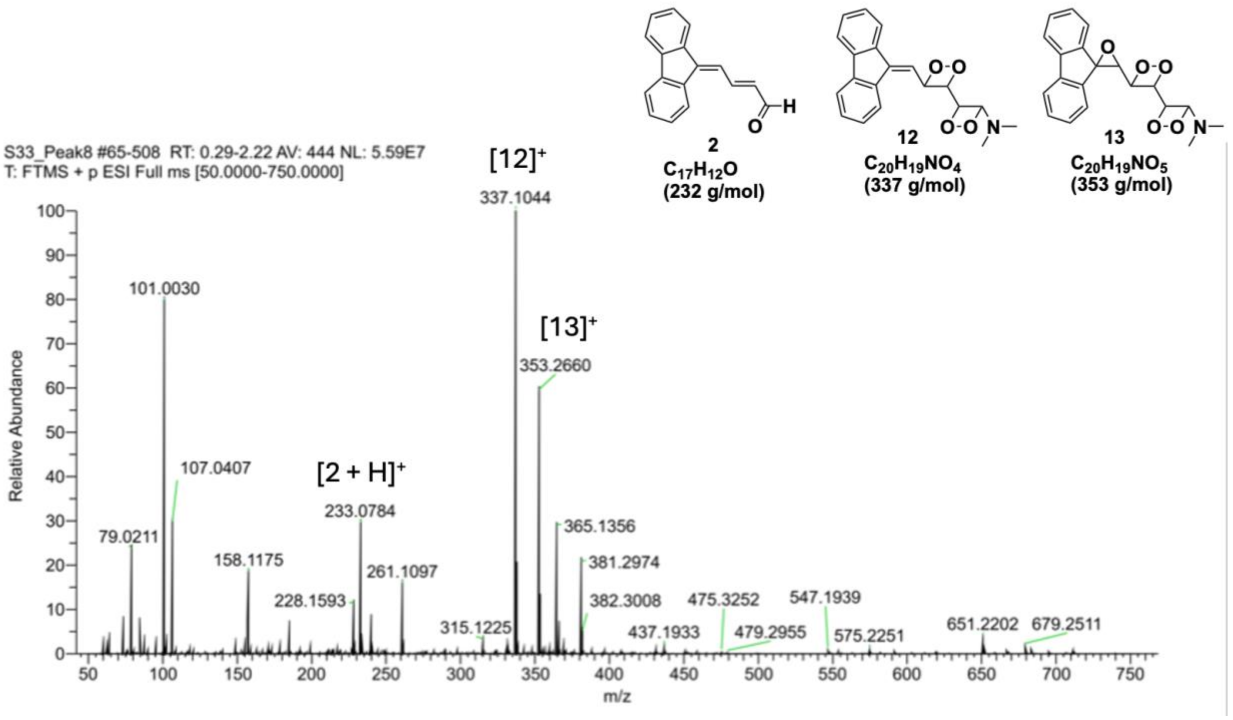


**Figure S13. Mass spectrum from peak 8 of the HPLC separation in Figure S10.**

| Bacterial isolate | DB10-R (μM) | DB10-Y (μM) |
|---|---|---|
| *S. aureus* USA100 | 25 | 100 |
| *S. aureus* USA200 | 25 | 100 |
| *S. aureus* USA300 | 25 | 100 |
| *S. aureus* USA400 | 6.25 | 100 |
| *S. aureus* USA600 | 25 | 100 |
| *S. capitis* | 25 | 100 |
| *S. epidermidis* | 25 | 100 |
| *S. chromogenes* | 25 | 100 |
| *S. lugdunensis* | 25 | 25 |
| *S. cohnii* | 50 | 200 |
| *S. warneri* | >200 | >200 |
| *B. subtilis* | 50 | 100 |
| *S. pyogenes* | 12.5 | 25 |
| *S. agalactiae* | 6.25 | 100 |
| *E. faecalis* | 25 | 200 |
| *P. aeruginosa* | >200 | >200 |
| *E. coli* | >200 | >200 |

**Table S1**: **DB33-R and DB33-Y are active against a variety of bacterial species.** MICs of DB33-R and DB33-Y against the same panel of bacterial strains and species previously tested for DB10-R and DB10-Y.

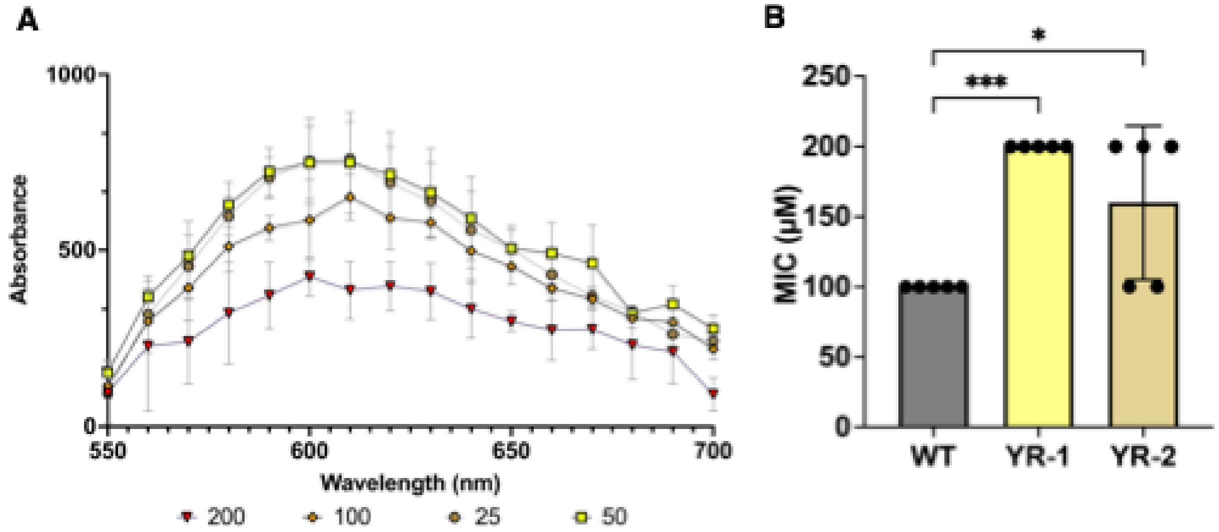


**Figure S14: Intercalation of DB33-Y into DNA.** (A) Like DB10-Y, DB33-Y intercalates into DNA, as shown by its ability to displace ethidium bromide (EtBr) from DNA, as measured by the drop in fluorescence. EtBr, DNA, and various concentrations of DB33-Y were incubated for 30 minutes in the dark before EtBr fluorescence spectra were measured (excitation at 525 nm). DB33-Y, EtBr, and DNA alone exhibited no intrinsic fluorescence. Data are presented as mean ± SD from three biological replicates. (B) DB33-Y exhibits reduced activity against *clpX* mutant strains YR-1 and YR-2, which were originally selected through enhanced resistance to DB10-Y.

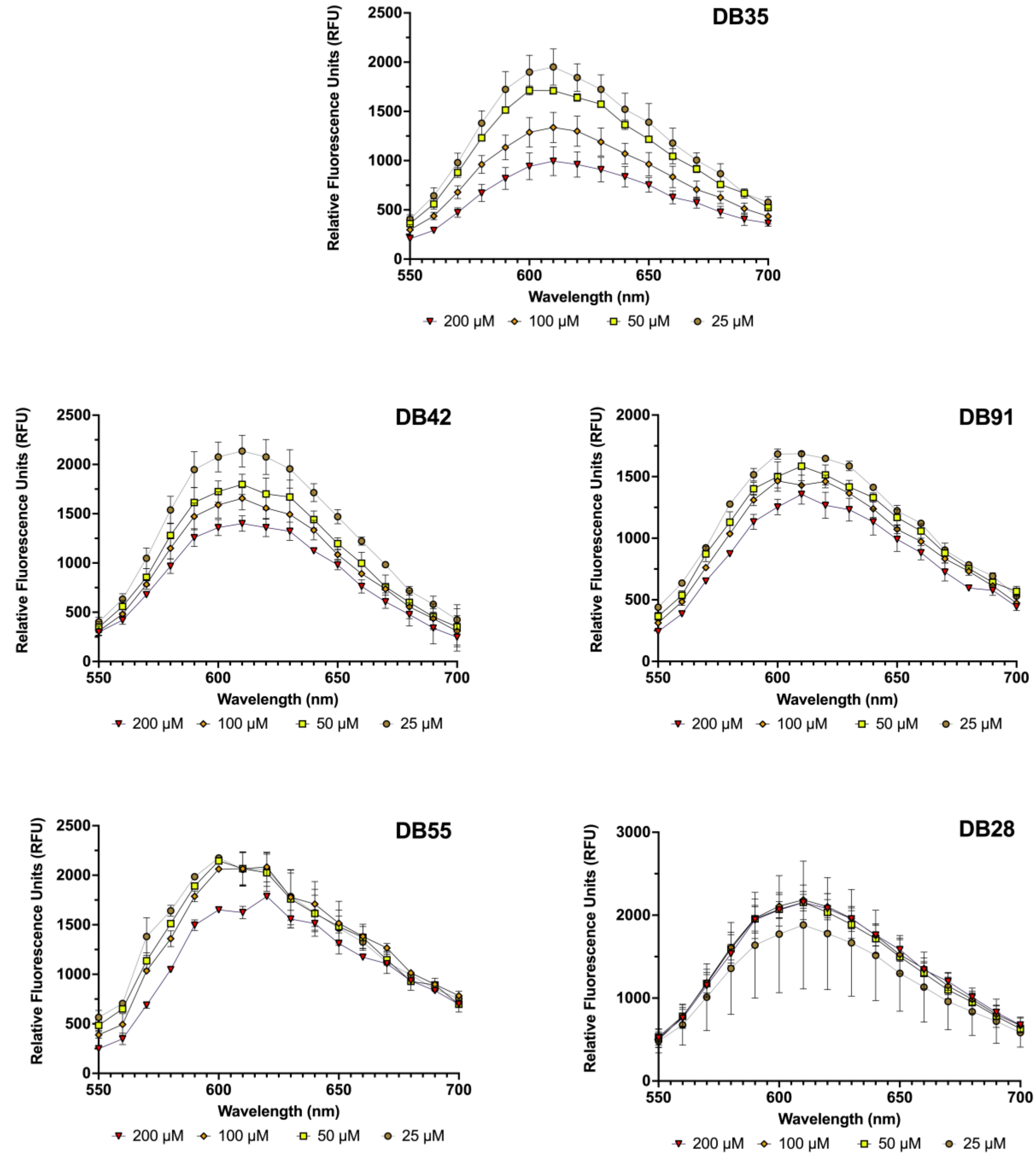


**Figure S15: Most DB10 analogs tested are DNA intercalators.** EtBr, DNA and various concentrations of analogs were incubated together for 30 minutes in the dark before the fluorescence spectra of EtBr was read (excitation 525 nm). DB35 and DB42 displaced EtBr at approximately the same rate as DB33-Y, DB91 and DB55 only slightly displaced EtBr at 200 µM. DB28 did not displace EtBr.

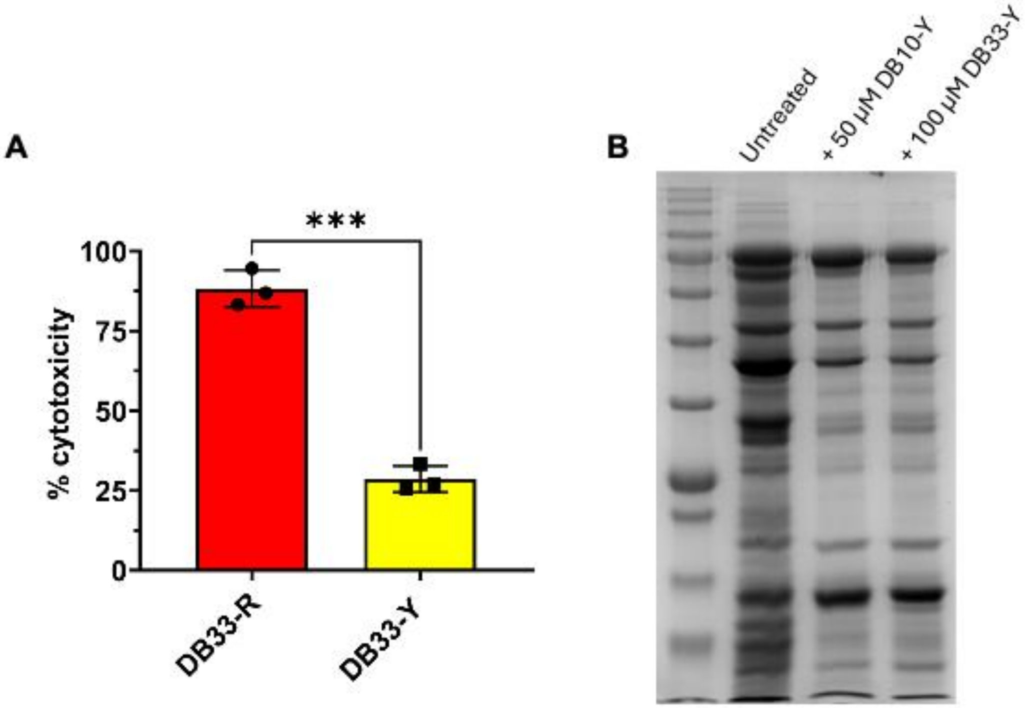


**Figure S16: DB33-Y inhibits intracellular *S. aureus* replication at non-cytotoxic concentrations and alters the bacterial secretome.** (**A**) Cytotoxicity of DB33 at 200 µM in RAW264.7 macrophages after 24h of exposure. Cytotoxicity was quantified using the LDH release kit, and used according to the manufacturer's instructions. Data are shown as the mean ± SD of at least three independent experiments. ∗p ≤ 0.05, ∗∗p ≤ 0.01, ∗∗∗p ≤ 0.001 using a one-way ANOVA with Dunnett's multiple comparison. (**B**) Full coomassie brilliant Blue R-250 stained gel of the cropped gel depicted in Figure 7F.

**Table S2: Bacterial strains used in this study**

| Strain or Plasmid | Description | Source or reference |
|---|---|---|
| ***S. aureus*** | | |
| USA300 | USA300 LAC, cured of resistance plasmids | Lab stock |

| | | |
|---|---|---|
| RN4220 | $r_K^- m_K^+$; capable of accepting foreign DNA | Lab stock |
| USA100 | WT *S. aureus* USA100 strain | Lab stock |
| BK21203 | WT *S. aureus* USA200 strain, MN8 lineage | Lab stock |
| MW2 | WT *S. aureus* USA400 strain | Lab stock |
| USA600 | WT *S. aureus* USA600 strain | Lab stock |
| USA300 *recA*::tn | WT *S. aureus* USA300 with a transposon mutation of *recA* | NTML library |
| USA300 *rexA*::tn | WT *S. aureus* USA300 with a transposon mutation of *rexA* | NTML library |
| USA300 *rexB*::tn | WT *S. aureus* USA300 with a transposon mutation of *rexB* | NTML library |
| USA300 *clpB*::tn | WT *S. aureus* USA300 with a transposon mutation of *clpB* | NTML library |
| USA300 *clpC*::tn | WT *S. aureus* USA300 with a transposon mutation of *clpC* | NTML library |
| USA300 *clpP*::tn | WT *S. aureus* USA300 with a transposon mutation of *clpP* | NTML library |
| USA300 *clpL*::tn | WT *S. aureus* USA300 with a transposon mutation of *clpL* | NTML library |
| USA300 p$clpX_{YR-1}$ | WT *S. aureus* USA300 carrying the mutant *clpX* gene (Codon 400 SNP) from USA300 on plasmid pALC2073 | This study |

| | | |
|---|---|---|
| USA300 p*clpX*$_{YR-2}$ | WT *S. aureus* USA300 carrying the mutant *clpX* gene (-12delC) gene from USA300 on plasmid pALC2073 | This study |
| **Other Staphylococcal species** | | |
| *S. epidermidis* Mach 1457 | Competent clinical strain | Lab stock |
| *S. lugdunensis* M23590 (HM-141) | Human skin isolate | ATCC type strain |
| *S. chromogenes* ATCC 43764 | From the original ATCC stock culture, this isolate possesses a white, mucoid colony | ATCC type strain |
| *S. capitis* ATCC 35661 | Human skin isolate | ATCC type strain |
| *S. warneri* ATCC 27836 | Human skin isolate | ATCC type strain |
| *S. cohnii* ATCC 29973 | Human skin isolate | ATCC type strain |
| **Other Gram-positive bacteria** | | |
| *M. luteus* ATCC 4698 | Human nasal secretion isolate | ATCC type strain |
| *B. subtilis* 3A1T | Wild-type isolate | Bacillus Genetic Stock Center |
| *S. pyogenes* MGAS8232 | Isolated from a patient with acute rheumatic fever. | J. McCormick |
| *S. agalactiae* A909 | Isolated from a septic human neonate | ATCC type strain |
| *E. faecalis* ATCC 33186 | Strain CN478; historical urine isolate | ATCC type strain |
| **Gram-negative bacteria** | | |

| | | |
|---|---|---|
| *P. aeruginosa* PAO1 | Wildtype strain | K. Poole |
| *E. coli* DH5α | F⁻ ϕ80d*lacZ*ΔM15 *recA1* *endA1* *gyrA96* *thi-1* *hsdR17*($r_K^-$ $m_K^-$) *supE44* *relA1* *deoR* Δ(*lacZYA-argF*)*U169 phoA* | Promega |
| **Plasmids** | | |
| pALC2073 | *E. coli - S. aureus* shuttle vector. $Amp^R$ in *E. coli*, $Cm^R$ in *S. aureus* | *82* |

**Table S3: Oligonucleotides used in this study**

| Name | Sequence | Description |
|---|---|---|
| *clpX*-F | TATATA GGTACC ATCTGCTACTATTCTTTAAGC | Forward primer to amplify *clpX* operon from *S. aureus* USA300 LAC |
| *clpX*-R | TATATA GAGCTC GATTGGAGCTTTTCACTTT | Reverse primer to amplify *clpX* operon from *S. aureus* USA300 LAC |
